# Rilmenidine mimics caloric restriction via the nischarin I1-imidazoline receptor to extend lifespan in *C. elegans*

**DOI:** 10.1101/2021.10.13.464294

**Authors:** Dominic F. Bennett, Anita Goyala, Cyril Statzer, Charles W. Beckett, Alexander Tyshkovskiy, Vadim N. Gladyshev, Collin Y. Ewald, João Pedro de Magalhães

## Abstract

Caloric restriction increases lifespan across species and has health benefits in humans. Because complying with a low-calorie diet is challenging, here we investigated pharmacological interventions mimicking the benefits of caloric restriction. Searching for compounds that elicit a similar gene expression signature to caloric restriction, we identified rilmenidine, an I1-imidazoline receptor agonist and prescription medication for the treatment of hypertension. We then show that treating *C. elegans* with rilmenidine at young and older ages increases lifespan. We also demonstrate that the stress-resilience, healthspan, and lifespan benefits upon rilmenidine treatment in worms are mediated by the I1-imidazoline receptor *nish-1*, implicating this receptor as a potential longevity target. Furthermore, we show that rilmenidine treatment increased ERK phosphorylation via NISH-1. Consistent with the shared caloric-restriction-mimicking gene signature, supplementing rilmenidine to caloric restricted *C. elegans*, genetic reduction of TORC1 function, or rapamycin treatment did not further increase lifespan. The rilmenidine-induced longevity required the transcription factors FOXO/DAF-16 and NRF1,2,3/SKN-1, both important for caloric restriction-mediated longevity. Furthermore, we find that autophagy, but not AMPK signaling, was needed for rilmenidine-induced longevity. Lastly, we find that treating mice with rilmenidine showed transcriptional changes in liver and kidney similar to caloric restriction. Overall, our findings reveal rilmenidine as a caloric restriction mimetic and as a novel geroprotective compound.

## Introduction

Individuals over 65 are now the fastest-growing demographic group worldwide, a fact that emblematizes the global aging population (Jarzebski et al. 2021). Unfortunately, at present, with age comes age-related chronic disease and death (Fontana et al. 2014), and as such, the estimated benefits of delaying aging, even if the effect is rather small, are immense (Farrelly 2010; Goldman et al. 2013; de Magalhães et al. 2012). A large body of evidence has demonstrated that the aging rate can be markedly slowed in model organisms. So far, caloric restriction (CR) is the most robust anti-aging intervention (Liang et al. 2018), and CR promotes longevity across species (Fontana et al. 2010). However, studies of CR in humans have had mixed results, low compliance, and many side effects (Most et al. 2017), meaning finding medications that can mimic the effect of caloric restriction is the most reasonable anti-aging target (Ingram & Roth 2015). However, only a few compounds have been identified to mimic the beneficial effects of CR.

Previously, we detailed a drug repositioning method in which we identified potential caloric restriction mimetics (CRMs), yielding biologically relevant results (Calvert et al. 2016). We did this by comparing drug-gene signatures to the *in-vitro* transcriptome of caloric restriction, looking for drugs with an overlapping profile. One drug identified during this work was the imidazoline receptor agonist allantoin, which was further confirmed to extend lifespan in *C.elegans* (Admasu et al. 2018). This has highlighted the potential of imidazoline agonists as caloric restriction mimetics. Allantoin, however, does not display oral bioavailability, obviating its use in humans (Kahn & Nolan 2000). We hypothesized, therefore, that a more potent and specific agonist of the imidazoline receptor, the widely-prescribed, oral antihypertensive rilmenidine (Dardonville & Rozas 2004), would have a similar, if not better, longevity effect in *C.elegans* but with more translatability to humans in the future.

Indeed, rilmenidine elicits a similar transcriptional profile to CR (Calvert et al. 2016). In another study, we also identified rilmenidine to reprogram human cell transcription profiles to a more youthful state (Statzer et al. 2021). Additionally, we identified rilmenidine by searching for gene expression signatures of longevity in mammals and subsequently have shown that the gene expression profile of liver from mice subjected to a diet containing rilmenidine for 1 month displays a strong positive association with the CR transcriptome profiles (Tyshkovskiy et al. 2019). Yet, whether rilmenidine acts as a CR mimetic to promote healthy aging is unknown.

In this work, we first validated the geroprotective properties of rilmenidine in *C.elegans* employing lifespan, healthspan, and stress assays. Rilmenidine extended the lifespan of *C.elegans* when commenced in early adulthood and also in aged animals. It did this via the nematode ortholog of the I1-imidazoline receptor (I1R/nischarin/IRAS), *f13e9.1/nish-1,* and was dependent upon increased autophagy and manipulation of three CR nexuses (DAF-16/FOXO, SKN-1/NRF, and mTOR) (Blackwell et al. 2015; Blackwell et al. 2019). Lifespan extension was not possible in genetic models of caloric restriction (*eat-2*), suggesting CR mimicry. Lastly, rilmenidine improved thermotolerance and attenuated PolyQ aggregation. This data demands further clinical investigation into the geroprotective effect of rilmenidine.

## Results

### Rilmenidine improves survival in *Caenorhabditis elegans*

Previous computational studies predicted rilmenidine as a longevity drug and CR mimetic (Calvert et al. 2016; Tyshkovskiy et al. 2019). To test if rilmenidine is involved in lifespan regulation, we treated wild-type (WT) *C. elegans* with different concentrations of rilmenidine at the larval stage L4 and performed lifespan assays. We found that rilmenidine extended the lifespan of WT animals by ∼19% compared to DMSO-treated WT control, at an optimal concentration of 200 μM (**Figure 1A, B, Supplementary Table 1**). Thus, we show that rilmenidine is indeed a geroprotective drug promoting longevity.

**Figure 1.**
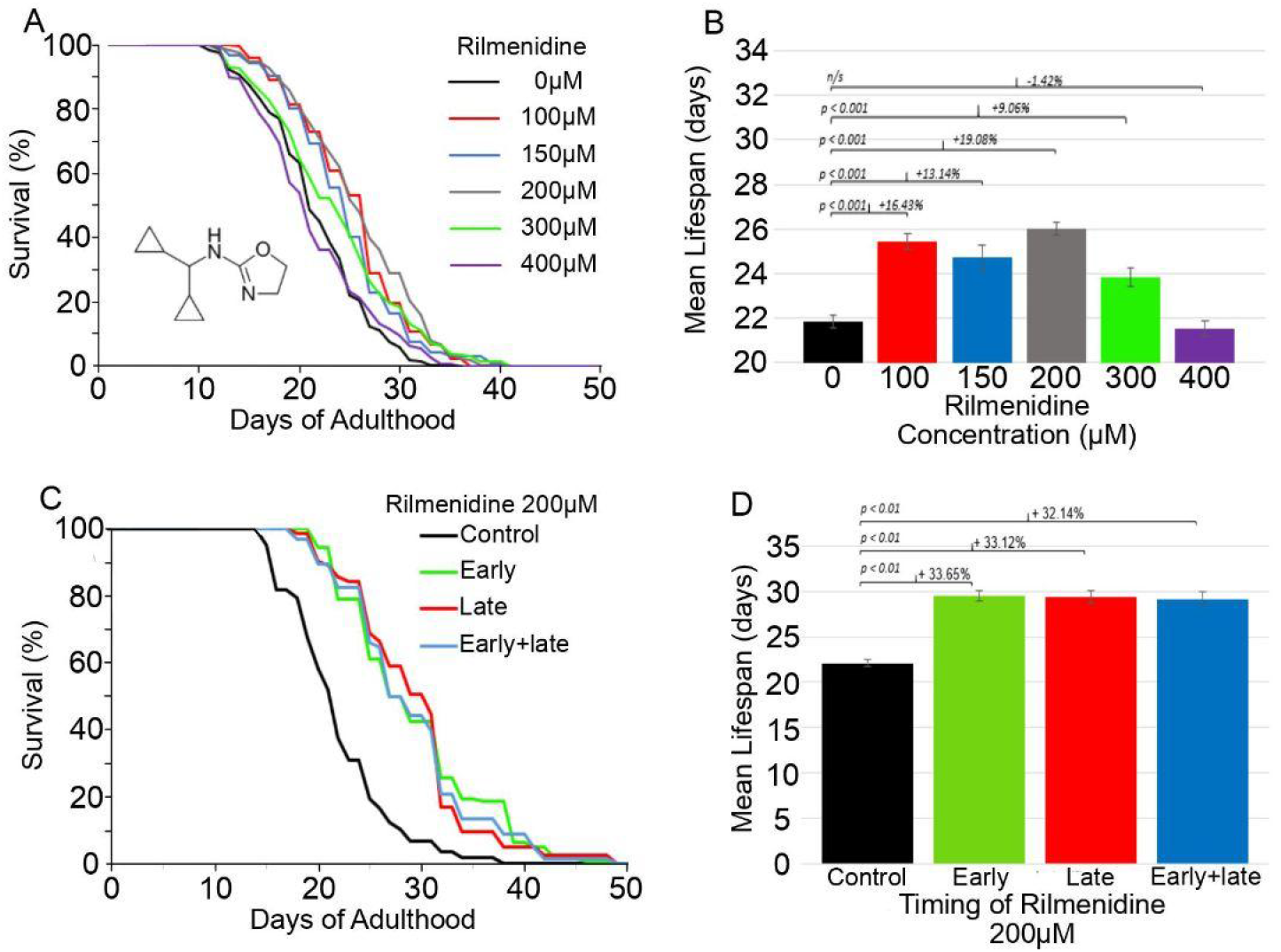
Improved survival of *C. elegans* treated with rilmenidine. **A)** Pooled survival curve showing significant lifespan extension in WT animals treated with rilmenidine at all concentrations, except 400 µM, compared to 1% DMSO vehicle control. **B)** Bar graph showing quantified lifespan data in terms of mean lifespan (days) for each rilmenidine concentration (100-400 µM) treated WT, compared to DMSO vehicle control; 200 µM concentration of rilmenidine provides maximum lifespan increase (19%) in WT. **C)** Pooled survival curve showing late-life treatment with rilmenidine (200 µM) at day 12 of adulthood increased lifespan in WT compared to DMSO vehicle control. **D)** Quantified data for lifespan assay showing percentage increase in mean lifespan of adult WT fed 200 µM rilmenidine at different times (day-1 or day-12 adulthood) compared to DMSO vehicle control. Error bars represent SEM; adjusted p-value was derived from log-rank test and Bonferroni correction. Kaplan Meir survival analysis was performed on pooled data from at least three independent trials. Quantitative data and statistical analyses for the representative experiments are included in Supplementary Table 1.

One goal is to administer geroprotective drugs at geriatric ages. However, not all longevity-promoting drugs work during old age. For instance, the geroprotective drug metformin, when administered at day 10 of *C. elegans’* adulthood or 20-months in mice, instead shortened lifespan and accelerated age-related pathologies (Espada et al. 2020; Zhu et al. 2021). Interestingly, we found that in worms rilmenidine prolonged the lifespan to the same extent (∼33%) whether exposure to the drug was initiated during youth, day 1 of adulthood, or when old, day 12 of adulthood (**Figure 1C, D, Supplementary Table 1**). However, this pro-longevity effect was not additive to lifespan extension exerted by early-life treatment, suggesting a possible cellular mechanism that upon activation lasts through adulthood and aging (**Figure 1C, D, Supplementary Table 1**). These results indicate that rilmenidine might be employed as a pharmacological intervention during old ages to extend lifespan.

### Identification of the conserved imidazoline receptor NISH-1 in *C. elegans*

Rilmenidine has been identified as a classical imidazoline type 1 receptor (I1R/IRAS/Nischarin) agonist in mammals (Zhang & Abdel-Rahman 2006). The amino-terminal sequence of Nischarin displays strong homology to a *C. elegans* protein encoded by the gene *f13e9.1* (**Figure 2A**) (Alahari et al. 2000). Our comparison of the *C. elegans f13e9.1* full-length protein sequence to the human orthologue IRAS protein sequence revealed high similarity in the following functional domains; PHOX (PX) domain, Leucine-rich repeats (LRRS), and coiled-coil (CC) domain (**Figure 2A**). The PHOX domain entails phosphoinositide-3-phosphate-binding capacity, likely enabling Nischarin to be anchored to the intracellular surface of the plasma membrane (Piletz 2003). The coiled-coil domain is the predicted imidazoline binding site (Sun et al. 2007) (**Figure 2A**). Thus, we renamed the *f13e9.1* gene to *nish-1* and will refer to it as *nish-1* throughout this study.

**Figure 2.**
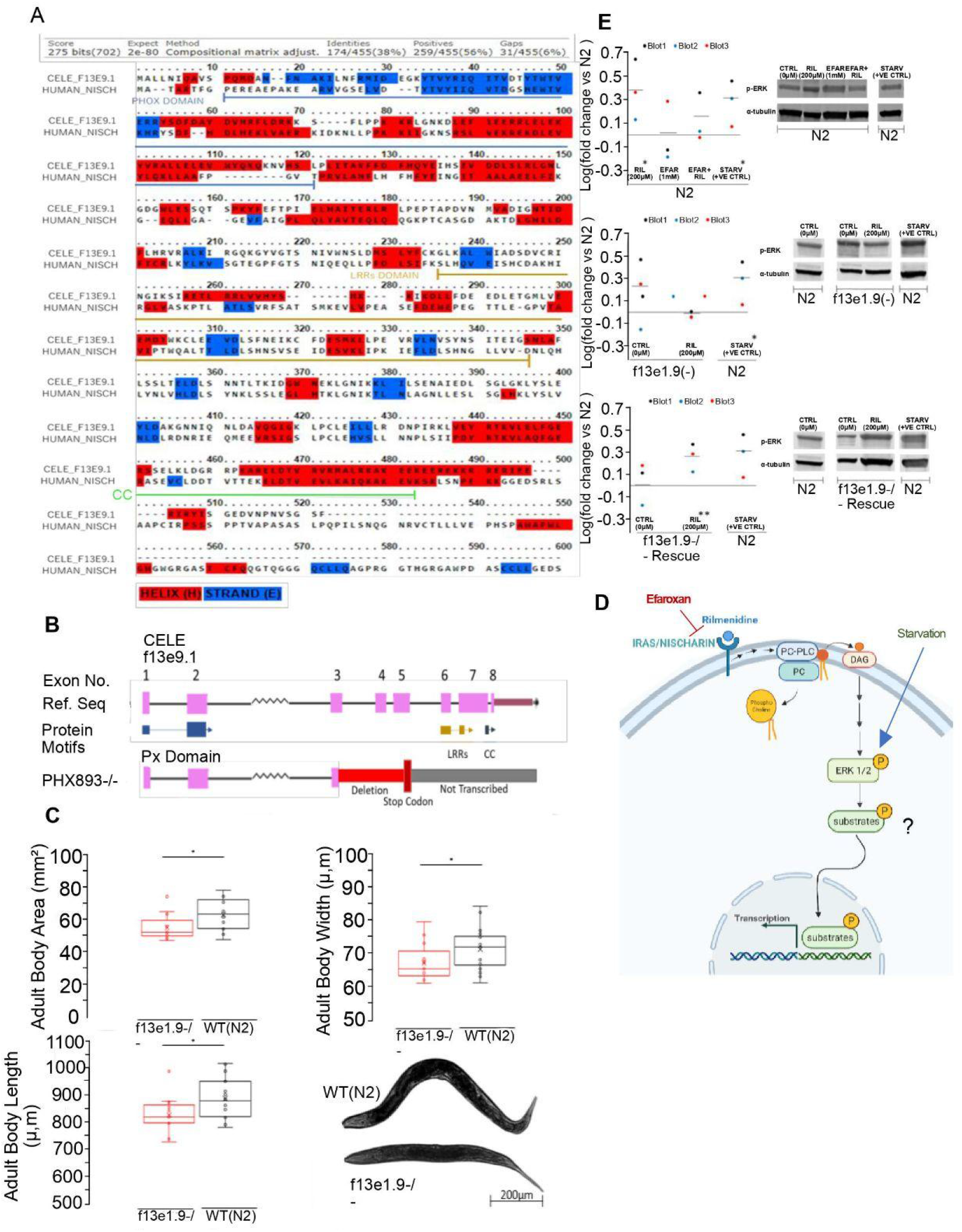
Characterization of NISH-1. **A)** BLASTp alignment of *C. elegans* NISH-1 protein sequence (CELE_F13E9.1 – UniProtKB-K8F807) against truncated human NISCHARIN isoform 3 (HUMAN_NISCH – UniProtKB-Q9Y2I1-3) revealing 38% sequence similarity, with conserved secondary structure elements like alpha helices shown in red and beta-strand in blue. Functional protein motifs like PHOX (Px) domains, LRRs, and coiled-coil domains are underlined. **B)** Schematic diagram illustrating exons of *nish-1* gene, alongside translated protein motifs and the deleted regions in the homozygous *nish-1* mutant. **C)** *nish-1* mutants exhibit reductions in body size compared to WT at day 1 of adulthood as shown by body area (defined as width x length). A two-tailed *t*-test was used for analysis; *p<0.05. Representative pictures of WT and *nish-1* mutant captured in brightfield at 10X objective on a Zeiss Axio Observer following paralysis in 20 mM tetramisole. Animal heads are left. Scale bar: 200 μm. **D)** Model illustrating the phosphorylation of ERK following interaction of rilmenidine to the NISH-1/IRAS receptor. **E)** Western blots showing MPK-1 phosphorylation in WT, *nish-1* mutants, or *nish-1* rescue transgenic animals, in response to 24h pharmacological intervention. α-tubulin was used as the loading control. Rilmenidine significantly increased MPK-1 phosphorylation *(%CV vs. FC = >1.5), however neither efaroxan alone nor rilmenidine and efaroxan in combination significantly increased MPK-1 phosphorylation n/s (%CV vs. FC = <1.5). Rilmenidine failed to reproducibly increase phosphorylation of MPK-1 in *nish-1* mutants. n/s (%CV vs. FC = <1.5). Transgenic rescue strains significantly increased MPK-1 phosphorylation upon rilmenidine treatment. **(%CV vs. FC = >2 in PHX945) and (%CV vs. FC =>1.5 in PHX946). Data is quantified as log-fold densitometric ratio change relative to α-tubulin. Data is then expressed as log fold change compared to DMSO vehicle control. Coefficient of variation (%CV), defined as the percent standard deviation: mean ratio; the significant difference in groups if % fold change is x1.5* greater or x2** than % CV.

To examine the role of *nish-1* as a functional ortholog of Nischarin that could mediate rilmenidine cell signaling, we generated a loss-of-function mutant *nish-1(syb767)* by CRISPR/Cas9 genome editing. A deletion of 873bp was initiated at the 3^rd^ exon and a stop codon was introduced to generate a potential null allele (see Materials and Methods for details). This resulted in truncated *nish-1* protein, containing only the PHOX domain but abolishing LRRs and the coiled-coil domain (**Figure 2B**). We validated the deletion in the *nish-1* mutant by genotyping (**Figure S 1A**).

Murine nischarin mutants are smaller than their wild-type counterparts (Crompton et al. 2017; Dong et al. 2017; Zhang et al. 2013). Therefore, for the functional characterization of the *nish-1* mutant, we first measured their body size. Day-1 adult *C. elegans nish-1* mutants were 5.44% smaller than WT (p-value <0.05; **Figure 2C**). To exclude slower development as the reason for size defects seen in *nish-1* mutants, we assessed the development rate of animals via the categorical staging of vulval development (Ludewig et al. 2017). Both WT and *nish-1* mutant developed at a similar rate and had reached full vulval development within 48h (**Figure S 1B**). Also, *nish-1* mutants did not exhibit any reproductive developmental abnormalities.

### Rilmenidine functions to activate ERK via *nish-1* in *C. elegans*

Previously, it has been shown that interaction of rilmenidine with the I1-imidazoline receptor leads to activation of phosphatidylcholine-specific phospholipase C (PC-PLC) and subsequent accumulation of the second messenger diacylglycerol (DAG) from phosphatidylcholine, with the release of phosphocholine. This leads to downstream activation of MAPK/ERK1/2, with phosphorylation of ERK1/2 being a standard read-out for the pathway (Zhang & Abdel-Rahman 2005) (**Figure 2D**). We utilized this well-characterized pathway read-out for testing if rilmenidine could activate the only orthologue of ERK, MPK-1 in *C. elegans*. The activation of MPK-1 was measured, by way of pMPK-1 to α-tubulin immunoreactivity, in response to 24 hours treatment with rilmenidine at varying concentrations (200 µM, 300 µM, and 400 µM) to establish a working concentration (**Figure S 1C**). Rilmenidine increased MPK-1 phosphorylation the strongest at the concentration of 200 µM. Since starvation also leads to phosphorylation of MPK-1, we used it as a positive control (You et al. 2006).

To establish whether this increase in activity was mediated via rilmenidine binding to an I1-imidazoline receptor, WT were co-incubated with rilmenidine (200 µM) and efaroxan (1 mM), an established selective I1-imidazoline receptor antagonist. 24 hours of efaroxan treatment alone did not significantly affect baseline MPK-1 activity compared to 1% DMSO vehicle-treated WT. However, efaroxan abolished MPK-1 activation by rilmenidine. This suggests rilmenidine treatment most likely interacts with an endogenous imidazoline binding site in *C. elegans* to elicit increases in MPK-1 activity (**Figure 2E**).

To further investigate whether the effect of rilmenidine on MPK-1 stimulation in *C. elegans* was mediated by the putative imidazoline ortholog NISH-1, we also measured MPK-1 phosphorylation in the *nish-1* mutant strain (**Figure 2E**). Rilmenidine failed to increase MPK-1 activity in *nish-1* mutants. (**Figure 2E**). In addition, in the transgenic strains wherein *nish-1* is rescued, the intensity of phosphorylation was higher in the presence of rilmenidine than in control, implying the necessity of *nish-1* for rilmenidine induced MPK-1 activation (**Figure 2E**).

### Rilmenidine requires *nish-1* for extending lifespan and improving healthspan

Next, we asked if rilmenidine extends lifespan through the *nish-1* receptor. We found that the lifespan-extending effects of rilmenidine were abolished when *nish-1* was deleted. Rilmenidine-treated *nish-1* mutants lived as long as WT and *nish-1* mutants without rilmenidine (**Figure 3A**). On the contrary, rescuing the *nish-1* receptor reinstated the increase in lifespan upon treatment with rilmenidine (**Figure 3B**). This demonstrates that *nish-1* plays a crucial role in relaying the signals downstream to increase lifespan.

**Figure 3.**
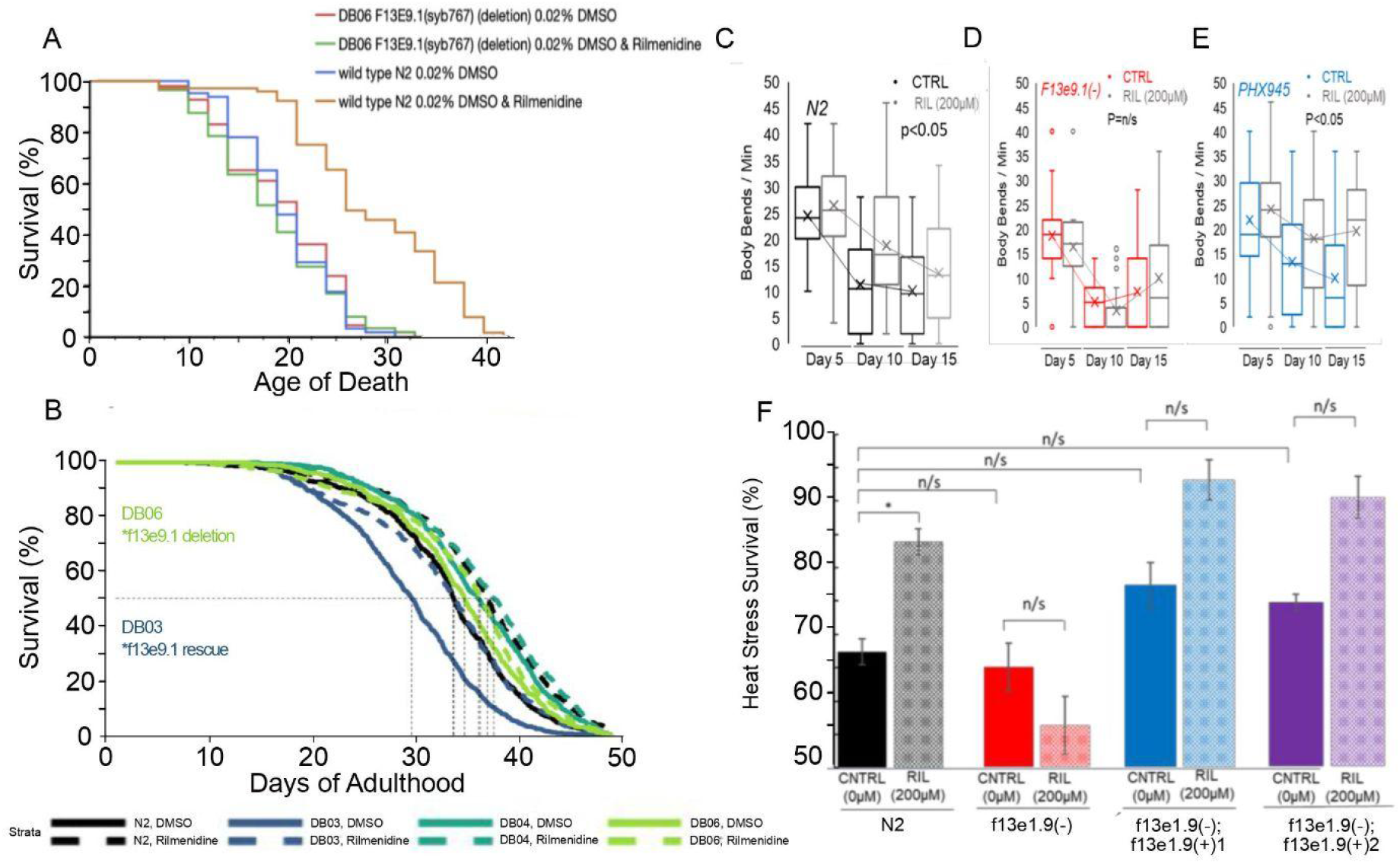
*nish-1* is required for an extended lifespan and better healthspan. **A)** Survival curve showing lifespan extension by rilmenidine in WT, but not in *nish-1* mutant (Statistics and raw data are in Supplementary Table 1). **B)** Rescue of *nish-1* enabled extension in lifespan by 200 µM rilmenidine in *nish-1* mutant. DB06 (=PHX893) is a *nish-1* knockout mutant and DB03 (=PHX945) and DB04 (=PHX946) are two rescue lines expressing wild-type *nish-1* gene copies. Quantitative data and statistical analyses for the representative experiments are included in Supplementary Table 1. **C)** Quantified data of body bends representing motility deterioration with age in WT **(C)**, *nish-1* mutants **(D)**, and two *nish-1* rescue mutants (PHX945 (=DB03) and PHX946 (=DB04)) **(E)** in the presence or absence of rilmenidine. Rilmenidine at 200 µM significantly reduced age-related motility deterioration in WT and *nish-1* rescue strains, but not in *nish-1* mutants. Data is represented as a pooled mean of 10 animals per time-point and condition/genotype repeated over three independent trials overlaid on a box plot representing quartiles. Error bars, upper: Q3 + 1.5*IQR; minimum: Q1 −1.5*IQR (adj. p<0.05; 2-way ANOVA and Tukey post hoc). **F)** Thermotolerance: Rilmenidine increases the percentage survival of WT exposed to 37°C for 3 hours and ∼20h recovery period, dependent on *nish-1*. Bar graph represents mean % survival ± S.E.M from three independent trials of at least 150 animals per strain and/or condition. *p<0.05 (one-way ANOVA with Tukey post hoc comparisons).

Further, we sought to explore if lifespan extension by rilmenidine is accompanied by better healthspan as well. Older animals reduce their movement with time, representing a marker of frailty (Hosono et al. 1980). Therefore, to investigate the healthspan of rilmenidine-treated WT animals, we measured swimming or thrashing assessed as body bends per minute. Rilmenidine showed higher thrashing rates with age in WT but not in *nish-1* mutants (**Figure 3C**). By contrast, *nish-1* mutants showed comparable numbers of body bends with or without exposure to rilmenidine, indicating beneficial effects of rilmenidine are conferred through the *nish-1* receptor (**Figure 3D**). The transgenic rescue of *nish-1* abrogated the age-related decline in the motility of *nish-1* mutants by increasing the number of body bends (**Figure 3E**). Thus, rilmenidine acts via the imidazoline receptor to improve healthspan as well.

### Rilmenidine requires *nish-1* to improve thermotolerance

Longevity is often associated with better stress survival (Zhou et al. 2011). Here, we determined the thermotolerance of animals exposed to rilmenidine. Rilmenidine treatment improved the thermotolerance at 37°C of WT animals but not *nish-1* mutants (**Figure 3F**). Transgenic rescuing the NISH-1 receptor in *nish-1* mutants restored rilmenidine’s resistance to heat stress (**Figure 3F**). This indicates that *nish-1* is central for rilmenidine to stimulate a conserved protective mechanism that may contribute to lifespan extension.

### Autophagy is required for extension in lifespan by rilmenidine

Rilmenidine has been shown to induce autophagy in SOD1 or TDP43 mutant mice models of amyotrophic lateral sclerosis (Perera et al. 2017; Perera et al. 2021). This warranted the examination of autophagy induction in *C. elegans* by rilmenidine. We used the mCherry::LGG-1 reporter strain to quantify autophagosome puncta in the posterior intestinal cells (Gosai et al. 2010; Li et al. 2014; Zhang et al. 2015). We found a significant increase in puncta formation by rilmenidine in a dose-dependent manner (**Figure 4A**). To determine whether the increase in autophagy is merely an association or has a causal effect on the longevity of rilmenidine, we performed lifespan assays and knocked down essential autophagy genes, *lgg-1* and *bec-1,* in WT exposed to rilmenidine or DMSO control. The lifespan extension conferred by rilmenidine was completely abrogated by impairment of autophagy (**Figure 4B, C, Figure S2, Supplementary Table 1**). This confirms that autophagy induction is a prerequisite for the increase in longevity by rilmenidine, similar to CR (Madeo et al. 2015).

**Figure 4.**
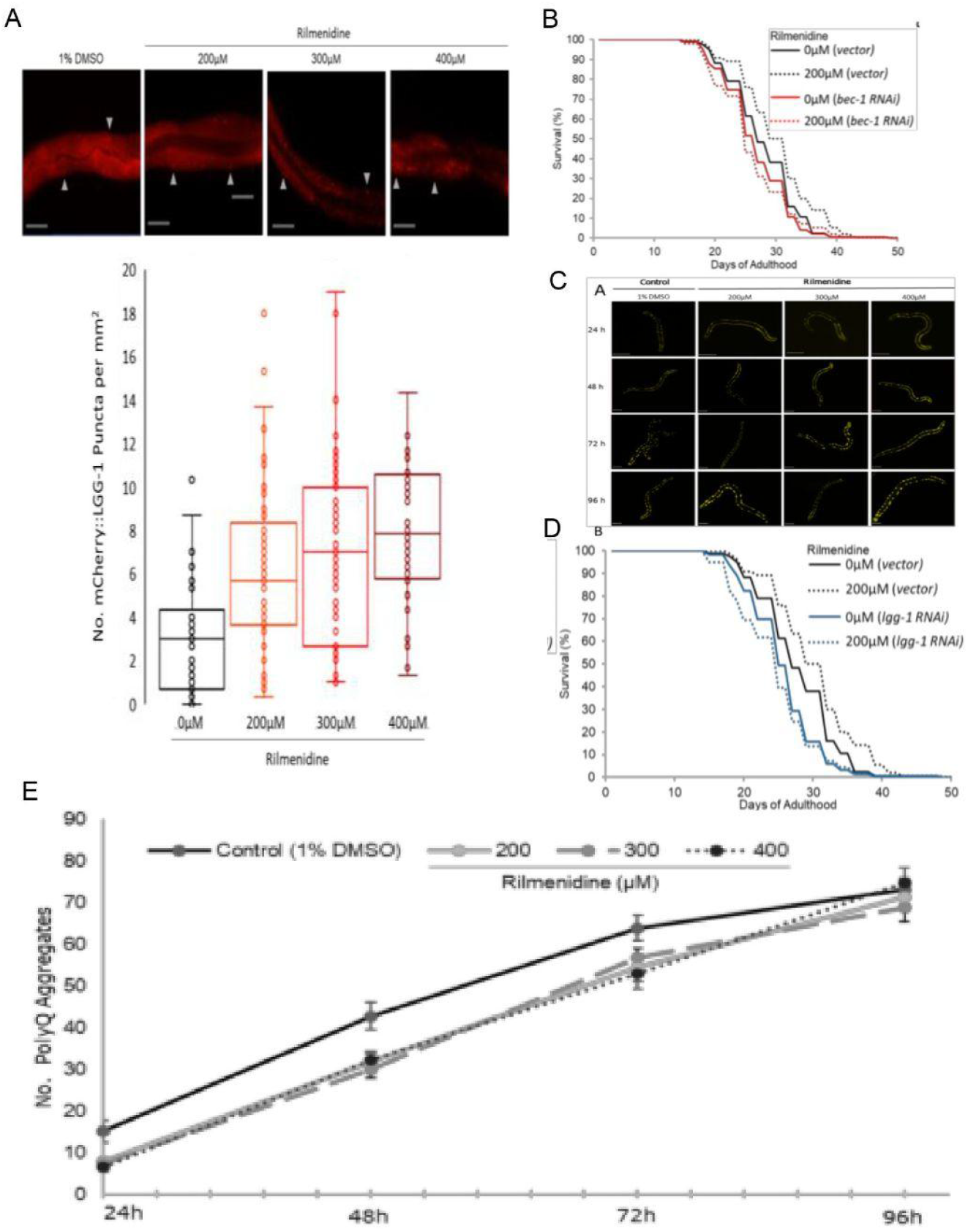
Induced autophagy by rilmenidine perturbed polyQ aggregation. **A)** Representative images of day 2 adult transgenic animals, expressing the intestinal specific autophagy reporter gene P*nhx-2::*mCherry::lgg-1 showing increased autophagy, when exposed to varying concentrations of rilmenidine for 24h compared to 1% DMSO vehicle. Arrows indicate autophagosome puncta formation. Scale bar = 20 μm. Graph shows the interquartile distribution of the mean number of mCherry::LGG-1 puncta in the posterior intestine of the animals in each condition. Error bars, upper: Q3 + 1.5*IQR; minimum: Q1 −1.5*IQR. **P<0.01, *P<0.05; one-way ANOVA followed by a Tukey’s post hoc test. **B,D)** Inhibition of autophagy abrogates the pro-longevity effect of rilmenidine as shown by survival curves of WT animals fed either RNAi bacteria expressing an empty vector (L4440), or *bec-1* **(B)** or *lgg-1* **(D)** dsRNA from day 1 adulthood in the presence or absence of 200 µM rilmenidine. Kaplan Meir survival analysis was performed on pooled data from at least three independent trials. Groups tested by Log-rank with Bonferroni correction; p<0.05. Quantitative data and statistical analyses for the representative experiments are included in Supplementary Table 1. **C,E)** Representative fluorescent micrographs of the transgenic strain repeated-measures ANOVA *p*<0.05.

### Rilmenidine attenuates poly-glutamine aggregates

One important hallmark of neurodegenerative diseases is the accumulation of protein aggregates. These should typically be degraded, perhaps via autophagy. Rilmenidine employs the process of autophagy as a therapeutic strategy reducing the soluble Huntington protein in the Huntington disease (HD) model in mice (Rose et al. 2010). Consistent with this finding in mice, we found that rilmenidine treatments significantly delayed the accumulation of polyQ40::YFP fusion protein aggregates compared to untreated WT (**Figure 4D**). Thus, rilmenidine may be considered for delaying the onset of neurodegenerative diseases like HD. Besides, this observation attests to our speculation of rilmenidine’s function as CRM, since DR too, mediates the reduction in the aggregation of polyQ in *C. elegans* (Matai et al. 2019).

### Rilmenidine acts as a CR mimetic

Rilmenidine shares the transcriptome profile with CR, as indicated by our meta-analysis studies (Calvert et al. 2016; Tyshkovskiy et al. 2019). Therefore, to discern the signaling pathway through which rilmenidine extends lifespan, we chose a genetic (*eat-2* mutant) model of CR in *C. elegans* (Lakowski & Hekimi 1998). Rilmenidine did not further increase the longevity of the *eat-2* mutants, suggesting that rilmenidine may function through the same genetic pathway (**Figure 5A, Supplementary Table 1**). CR is thought to work through the nutrient sensor, mTOR complex 1, mTORC1 to extend lifespan (Hansen et al. 2007; Statzer et al. 2020). Also, mTOR inhibition leads to autophagy induction, like rilmenidine (Hansen et al. 2007). To investigate the requirement of mTOR in lifespan increase by rilmenidine, we performed lifespan assays using a heterozygous mutant of *daf-15*, the raptor adaptor protein in mTORC1. We found that rilmenidine could not increase the longevity additively of the *daf-15(m81/+)* mutants (**Figure 5B, Figure S3, Supplementary Table 1**). Complementary, we inhibited mTOR pharmacologically by rapamycin. Rapamycin significantly increased the lifespan of WT animals. Co-treatment with rilmenidine and rapamycin failed to further increase the mean lifespan extension of WT animals treated either with rilmenidine or rapamycin individually (**Figure 5C, Figure S3, Supplementary Table 1**). This result strengthens the earlier observation that rilmenidine indeed acts through mTOR1 signaling for increasing the lifespan.

**Figure 5.**
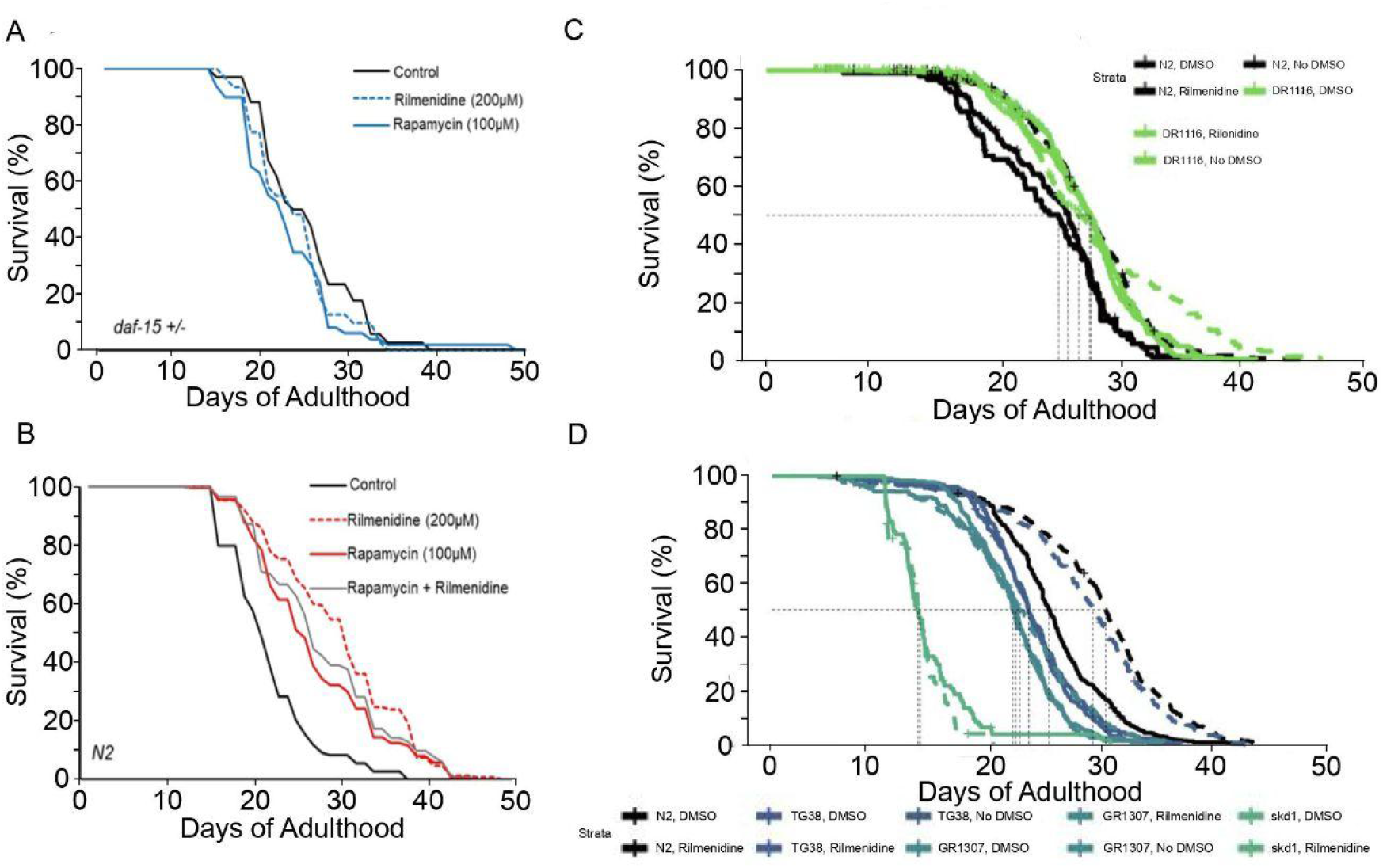
Effects of rilmenidine treatment on survival of CR-associated mutants. Survival curves showing the inability of rilmenidine to extend life span in **(A)** DR412 *daf-15(m81/+)* mutants**, (B)** rapamycin-treated WT, **(C)** DR1116 *eat-2(ad116)* mutants, and **(D)** GR1307 *daf-16(mgDf50)*, LD1057 *skn-1(tm3411)*, but TG38 *aak-2(gt33)* mutants. Raw data, quantitative data, additional trials, and statistical analyses for the representative experiments are included in Supplementary Table 1.

In *C.elegans*, mTOR inhibition by genetic or pharmacological intervention, like in mammals, leads to activation of key longevity transcription factors such as SKN-1/NRF and DAF-16/FOXO (Robida-Stubbs et al. 2012). Moreover, on solid media, *daf-16* is indispensable for DR-induced longevity (Greer et al. 2007). To explore whether rilmenidine induced longevity effects are DAF-16 and SKN-1 driven, we employed *skn-1(tm3411)* and *daf-16(mgDf50)* null mutants to perform lifespan assays. We found that rilmenidine did not significantly extend the lifespan of either *daf-16* or *skn-1* mutants, revealing a requirement of *daf-16* and *skn-1* for longevity benefits by rilmenidine (**Figure 5D, Supplementary Table 1**).

AMPK is another critical metabolic switch, which works antagonistically with the mTOR pathway and is activated by CRM drugs like metformin (Zhang et al. 2019; Onken & Driscoll 2010). Thus, we assessed the necessity of AMPK in the mediation of rilmenidine’s pro-longevity effects. We found that rilmenidine additively extended lifespan in *aak-2 (gt33)* mutants compared to WT, suggesting AMPK is not required for lifespan regulation by rilmenidine (**Figure 5D, Supplementary Table 1**).

All the above results show that rilmenidine functions through the imidazoline receptor, *nish-1*, which activates genetic pathways mediated by the established CR nexuses to extend the lifespan.

### Rilmenidine produces longevity-associated gene expression effects in mouse tissues

To test if the observed beneficial effects of rilmenidine are recapitulated in mammals, we performed RNA-seq analysis of liver and kidney samples from young UM-HET3 male mice subjected to rilmenidine for one month, together with the corresponding controls. We identified transcriptomic effects of the drug independently for each tissue and compared them with biomarkers of lifespan-extending interventions and aging, using a gene set enrichment analysis (GSEA) based approach (**Figure 6A**) (Tyshkovskiy et al. 2019). Gene expression signatures of lifespan extension reflect a liver response to individual interventions, such as caloric restriction (CR), rapamycin and mutations leading to growth hormone (GH) deficiency; gene expression changes shared by different longevity interventions; and genes, which expression is quantitatively associated with the effect on maximum and median lifespan (Tyshkovskiy et al. 2019). Aging signatures correspond to murine age-related gene expression changes in the liver and kidney, according to the Tabula Muris Consortium (Schaum et al. 2020).

**Figure 6.**
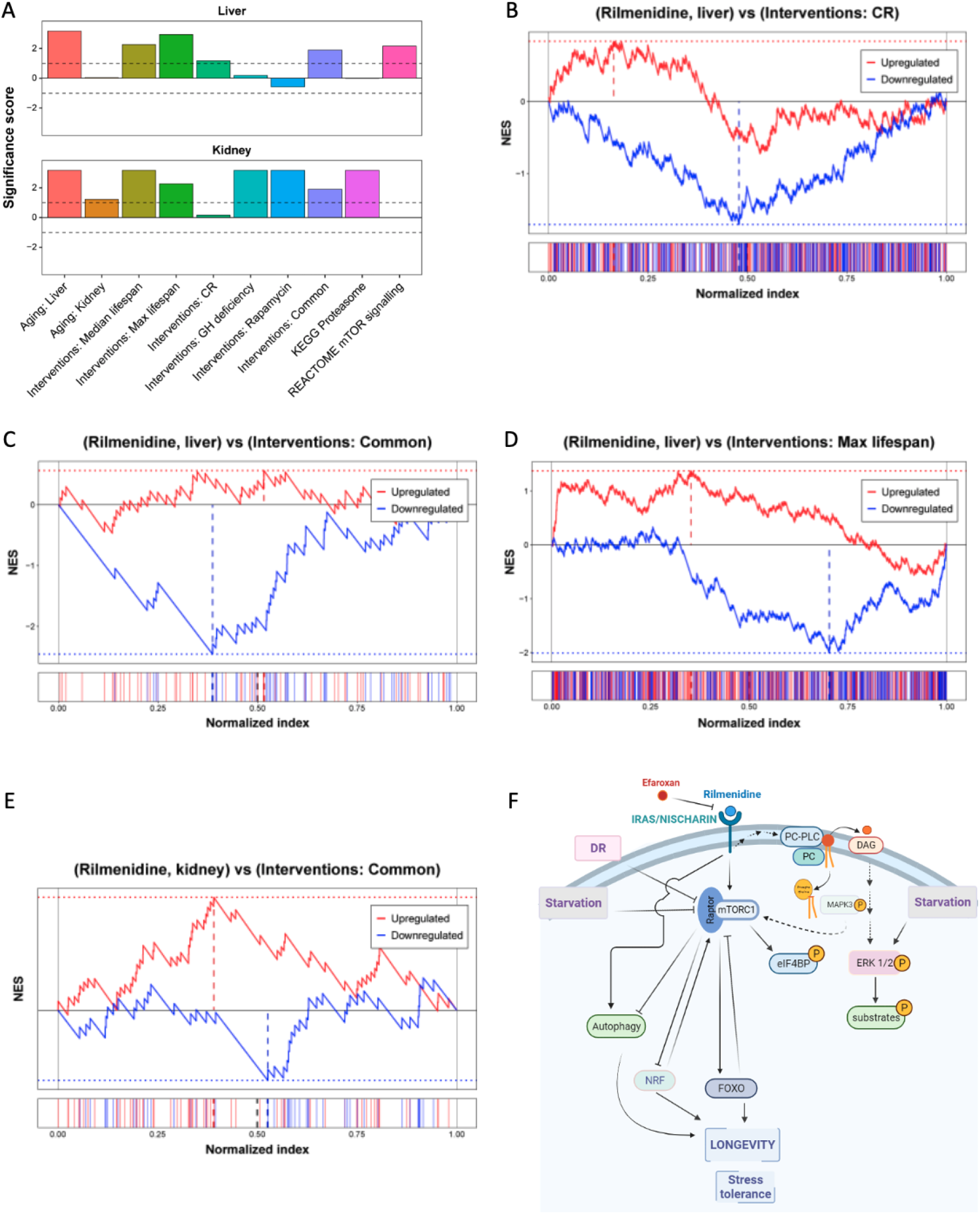
Rilmenidine treatment in mice induces gene expression changes associated with a lifespan-extending effect. **A)** Association of the liver (top) and kidney (bottom) responses to rilmenidine with signatures of aging (left), lifespan-extension (middle), and longevity-related intracellular processes (right). The significance score was calculated as log_10_(adjusted p-value) corrected by a sign of regulation. Dotted lines represent the FDR threshold of 0.1. **B-E)** GSEA enrichment plots with running normalized enrichment scores (top) and distributions (bottom) of selected up- (red) and downregulated (blue) gene signatures of lifespan-extending interventions among genes perturbed by rilmenidine in the liver **(B-D)** and kidney **(E)**. Perturbed genes were sorted based on the log_10_(p-value) of their differential expression between control and rilmenidine-treated samples corrected by a sign of regulation. The resulting index was divided by the number of genes in the dataset. Dotted lines represent NES calculated for up- (red) and downregulated (blue) gene signatures. **F)** Working model representing possible pro-longevity signaling by rilmenidine to extend life span in *C. elegans*. NES: Normalized enrichment score; CR: Caloric restriction; GH: Growth hormone; mTOR: mammalian target of rapamycin.

Interestingly, changes induced by rilmenidine in both liver and kidney were positively associated with the majority of longevity signatures, including biomarkers of median and maximum lifespan extension (adjusted p-value < 2.10^-3^ for both liver and kidney), CR (adjusted p-value = 0.066 for liver), rapamycin (adjusted p-value = 6.7.10^-4^ for kidney) and common signatures of longevity interventions (adjusted p-value < 0.013 for both liver and kidney). In other words, genes up- and downregulated in long-lived mice were also changed in the same direction after rilmenidine administration (**Figure 6B-E**). Remarkably, we also observed a significant positive association of rilmenidine effect with biomarkers of aging (median adjusted p-value = 0.03), which may be partly due to the age-related downregulation of insulin and IGF-1 signaling (López-Otín et al. 2013).

To understand if rilmenidine in mice regulates cellular functions similar to those identified in *C. elegans*, we also performed functional GSEA. This method allowed us to test if genes involved in certain pathways, based on REACTOME, KEGG and GO BP annotation, were enriched among those affected by the compound. We observed significant enrichment of genes involved in mTOR signaling and proteolysis among genes upregulated by rilmenidine in murine liver and kidney, respectively (adjusted p-value < 7.10^-3^) (**Figure 6A**). These results suggest that longevity-associated molecular mechanisms induced by rilmenidine in *C. elegans* are reproduced in mammals, pointing to a conserved geroprotective effect and downstream signaling (**Figure 6F**).

## Discussion

Challenges with humans’ long-term caloric restriction (CR) routines raise an unmet need to explore pharmacological interventions like calorie restriction mimetics (CRM). These may provide an alternative to CR, conferring the same longevity benefits without the challenges or side effects of low-calorie diets. Studies detailing the therapeutic efficacy of CRMs when administered at geriatric stages are rarely performed (Ingram & Roth 2015). Identifying drugs that can mirror the benefits of long-term calorie restriction even when the administration is initiated at late stages of life are of great interest.

Our study illustrates that rilmenidine acts as a CRM in *C. elegans*, improving survival, even when administered at older ages (commencing day-12 of adulthood). Rilmenidine targets conventional signaling nodes like mTOR, DAF-16/FOXO, SKN-1/NRF in *C. elegans* and activates stress response pathways, enhancing autophagy, attenuating polyglutamine aggregation, and improving thermotolerance. Rilmenidine acts via a novel mechanism wherein the newly identified imidazoline receptor, *nish-1* (previously *f13e9.1*) in *C. elegans*, mediates signaling to the genetic pathways relating to CR to extend lifespan. Although the imidazoline receptor signaling had been implicated to interact with alpha/beta-integrin or insulin-dependent PI3 kinase/Akt pro-survival pathways (Dontenwill et al. 2003), the connection of the imidazoline receptor to CR or longevity had not been studied. We showed that rilmenidine-induced lifespan and healthspan extension are dependent upon the imidazoline receptor, *nish-1*.

Many known CRMs, like metformin, rapamycin, or resveratrol, had been shown to act via AMPK activation, mTOR inhibition, and sirtuin activation, respectively (Shintani et al. 2018). Rilmenidine too can be viewed as a CRM as it regulates the intracellular signaling systems like the mTOR pathway in *C. elegans*. Indeed, rilmenidine induces the recycling process of autophagy, which is considered one of the characteristic features of CRMs and mTORC1 signaling (Chung & Chung 2019; Mariño et al. 2014). In mouse models, rilmenidine induced similar changes in mTOR regulation and autophagy activation in liver tissue (Figure 6). Since mTOR signaling and AMPK are two key players in regulating autophagy, and AMPK is not involved in lifespan extension by rilmenidine, rilmenidine probably induced autophagy by inhibition of inhibition mTOR signaling. The exact signal transduction pathway from the interaction of rilmenidine with *nish-1* to the inhibition of mTOR might be worth addressing in future studies. Rilmenidine has been reported to enhance the phosphorylation of ERK/MAPK via the phosphatidylcholine-specific phospholipase C pathway (Zhang & Abdel-Rahman 2005). We speculate, therefore, that ERK activation may inhibit the mTOR complex. In fact, mTOR inhibition upregulates phospho-ERK signaling through p70S6K in a feedback loop (Sunayama et al. 2010; Alonso et al. 2016). This might explain the upregulation of phospho-ERK upon rilmenidine binding to nischarin-1/IRAS. In any case, questions remain as to what signal transduction components are implicated by rilmenidine to induce autophagy.

By modulating autophagic flux, CRMs protect against neurotoxic protein aggregates (Yang & Zhang 2020; Sanchis et al. 2019). Previous studies reported autophagy to be a major checkpoint for the protein aggregates in many neurodegenerative polyglutamine diseases such as spinocerebellar ataxias, Huntington disease, etc, among model organisms (Ravikumar et al. 2002; Ravikumar et al. 2004). For instance, pharmacologically inducing autophagy using the CRM rapamycin encouraged the degradation of ataxin-3 aggregates (Ravikumar et al. 2004; Berger et al. 2006). Similarly, rilmenidine was proven effective for removing polyQ aggregates in mice models of Huntington disease and amyotrophic lateral sclerosis by upregulating autophagy (Rose et al. 2010; Perera et al. 2018). We further showed rilmenidine to be beneficial against the polyQ aggregates in *C. elegans*. Additionally, we observed upregulation of proteolysis in the rilmenidine treated kidney tissue in mice, contributing to increased clearance of the aggregated proteins.

Cytoprotective autophagy is also activated by heat shock response (Kumsta et al. 2017). Furthermore, the heat shock response helps to curb proteotoxicity due to aggregation (Hsu et al. 2003). We found that rilmenidine improved thermotolerance in *C. elegans.* These observations indicate that autophagy induction by rilmenidine is an important stress evasion mechanism, which supports proteostasis, possibly through regulating the proteasome pathway and heat stress response and thus, extending lifespan.

Alongside being a clinically-approved anti-hypertensive drug, rilmenidine improves plasma lipid and blood glucose in patients with hypertension and metabolic syndrome (Luca et al. 2000; Reid 2001). Given its contribution towards lowering blood glucose and increasing insulin sensitivity, rilmenidine may have an anti-diabetic function. Since I1R and I3R do not share similar ligand binding sites and I3R is a regulator of insulin secretion, the potential interaction of rilmenidine with the I3type imidazoline receptor might explain its effects on improved insulin resistance and, thus, glucose levels (Bousquet et al. 2020). As rilmenidine improves glucose tolerance, it can likely be put in the insulin sensitizer class of CRMs (Shintani et al. 2018, Ingram et al. 2006).

Since it is already in use, it becomes feasible for rilmenidine to be repositioned against insulin resistance, metabolic syndrome, and polyglutamine diseases. Moreover, conservation of findings between *C. elegans,* mice, human cell culture, such as induction of autophagy, points towards the more considerable potential for translating the longevity benefits to humans. In conclusion, rilmenidine is a new addition to the list of potential CRMs that could be viable therapeutic interventions administered late in life, and thus, warrants further examinations.

---

**Figure S1.**
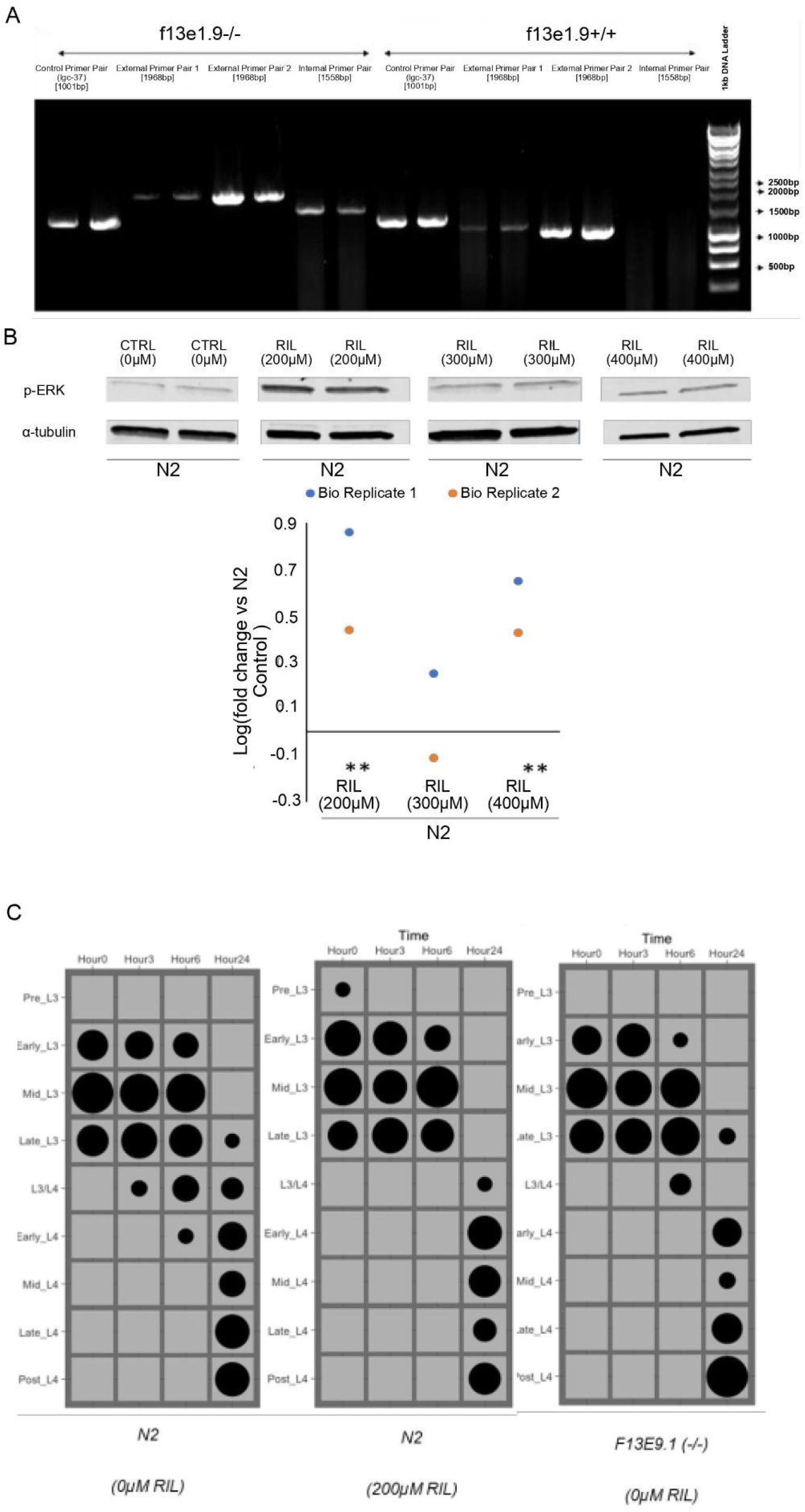
**A)** Endpoint single worm PCR genotyping of WT and *nish-1* deletion mutant following x4 outcrossing. Using oligonucleotides flanking the deletion site (external primer pairs) generated either a 1968 bp or a 1936 bp amplicon depending on oligonucleotide pair in WT or a correspondingly 873 bp reduced amplicon size in the genomic DNA of homozygous *nish-1* mutants. Likewise, an oligonucleotide pair targeted to sequences within the CRISPR deletion site produced a 1558 bp amplicon in WT but no amplicon in homozygous *nish-1* mutants. Control amplifications were conducted on the *lgc-37* gene coded on chromosome III. Primer sequences are in Supplementary Table 2. **B)** Effect of different concentrations on rilmenidine-induced MPK-1 phosphorylation as measured by Western Blotting. Phosphorylation of MPK-1 was assessed in WT in response to 24h pharmacological intervention. Data is then expressed as log fold change compared to 1% DMSO vehicle control. Rilmenidine at 200µM and 400µM significantly increased MPK-1 phosphorylation Coefficient of variation (%CV), defined as the percent standard deviation: mean ratio; significant difference in groups if % fold change is x1.5* greater or x2** than % CV. **(%CV vs. FC = >2).. **C)** *nish-1* mutants and rilmenidine-fed WT do not exhibit delayed vulval development from the L3 stage. In the chart, developmental stages are listed on the Y-axis, and the number of hours after a 24h exposure to food from L1 is plotted on the x-axis. The areas of the circles in the chart reflect the percentage of the population at each stage of development; n≥30 from 3 independent pooled trials for each time point. Statistical difference established by way of 2-way ANOVA wherein a mean development score could be ascertained through pre-L3 corresponding to a score of 0, Early L3 is 1 and Mid L3 a 3 etc. No groups displayed any significant difference in development rate.

**Figure S2:**
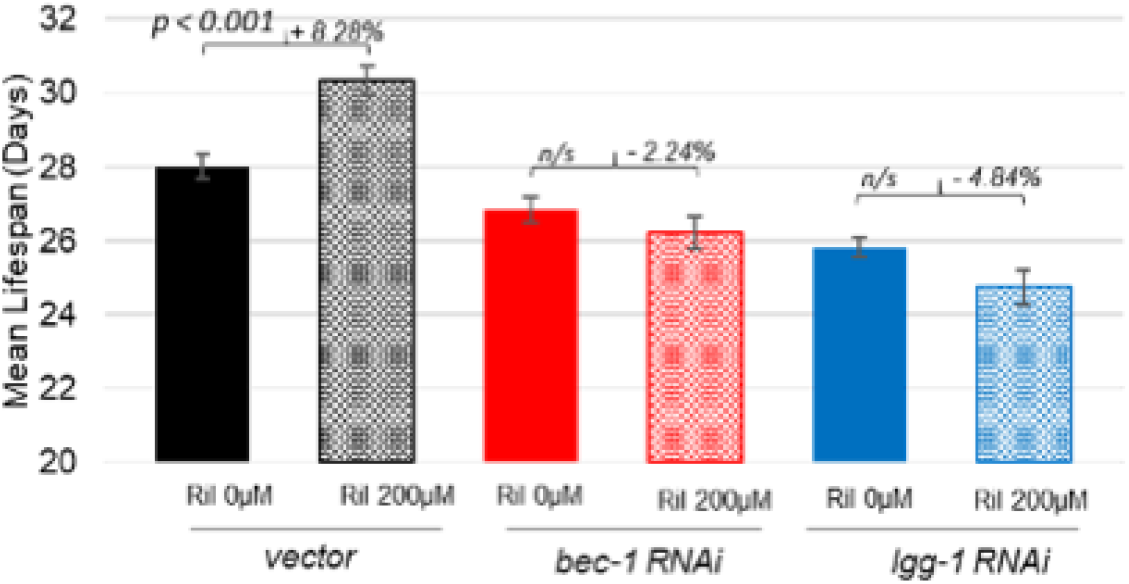
Related to figure 4B and 4C. Quantified data for mean lifespan (days) and % change in survival of WT upon knockdown of autophagy genes, *bec-1* and *lgg-1* alongside rilmenidine treatment. Rilmenidine significantly increased lifespan by 8.29% in WT animals fed HT115 *E.coli* expressing an empty vector (p<0.001), however, failed to significantly affect lifespan in animals fed HT115 *E.coli* expressing *bec-1* or *lgg-1* dsRNA; n/s = p>0.05. See Supplementary Table 1 for details.

**Figure S3:**
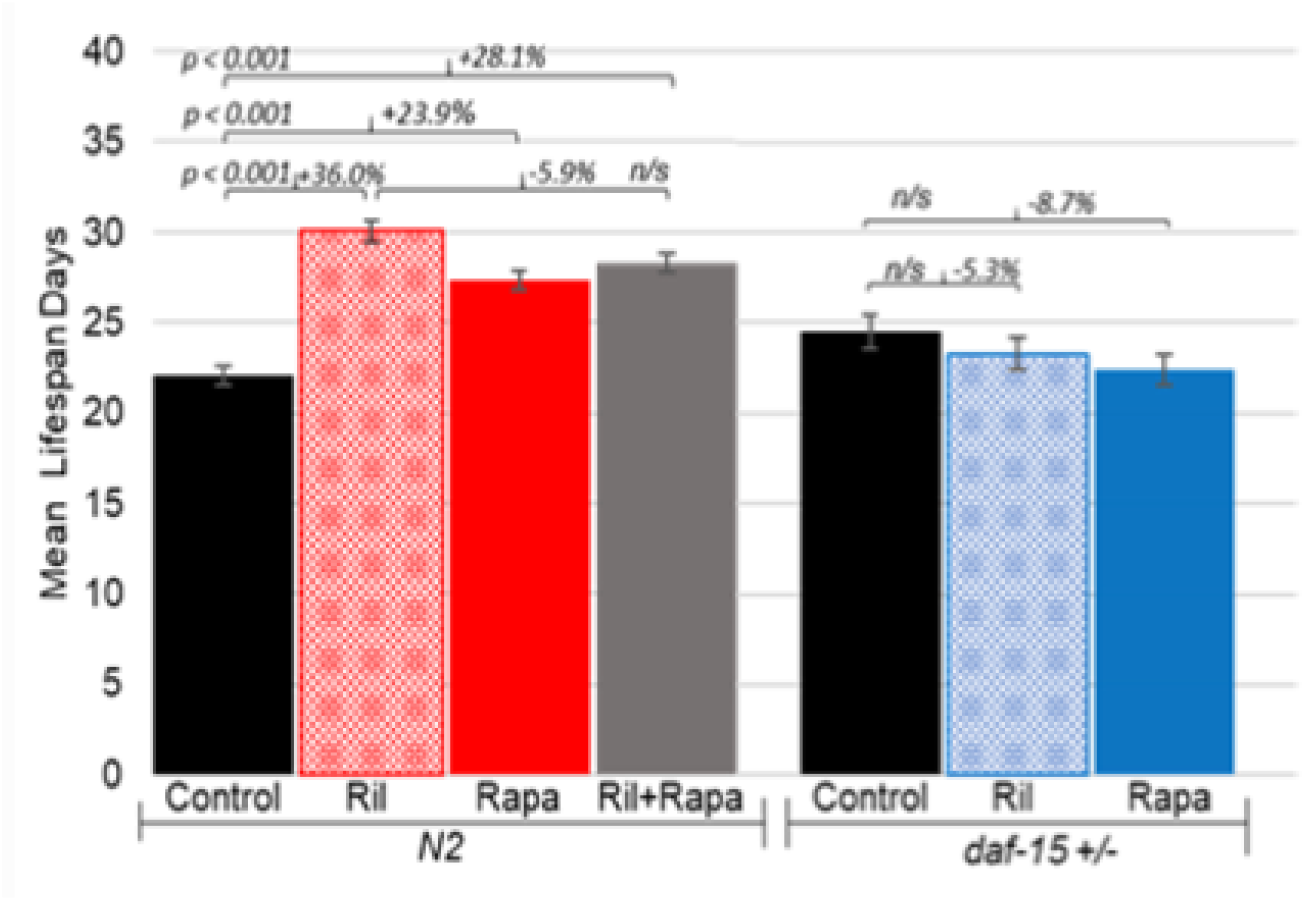
Related to figure 5B and 5C. Quantified data for mean lifespan (days) and % change in lifespan upon rilmenidine administration in *daf-15(m81/+)* mutants and rapamycin-treated WT. Error bars represent ± SEM; adjusted p-value was derived from log-rank test and Bonferroni correction. See Supplementary Table 1 for details.

**Supplementary Table 1: Lifespan summary and details.**

**Supplementary Table 2: Primer sequences.**

## Materials and Methods

### C. elegans Strains

All strains were maintained on OP50 *E.coli* seeded NGM agar supplemented with 50 IU/mL penicillin (Sigma #P4333), 50 µg/ml streptomycin (Sigma #P4333), and 50 IU/mL Nystatin (Sigma #1638) to prevent contamination (Stiernagle 2006). The strains used in the study are: WT (N2 Bristol), PHX893 (=DB06) *nish-1(syb767),* PHX117 *sybIs62* [*f13e9.1*::EGFP::3xFLAG, *unc-119*(+) + P*myo-2*::mCherry], PHX116 *sybIs62* [*f13e9.1*::EGFP::3xFLAG, *unc-119*(+) + P*myo-2*::mCherry], PHX945 (=DB03) and PHX946 (=DB04) are two independent integrated lines of *f13e9.1 (syb767)* IV; *sybIs62* [*f13e9.1*::EGFP::3xFLAG, *unc-119*(+) + P*myo-2*::mCherry], VK1093 *vkEx1093* [P*nhx-2*::mCherry::*lgg-1*], AM141 *rmIs133* [P*unc-54*::Q40::YFP], GR1307 *daf-16(mgDf50)* I; DA1116 *eat-2(ad1116)* II; LD1057 *skn-1(tm3411)* (Ewald et al. 2015) IV, DR412 *daf-15(m81)/unc-24(e138)* IV; and TG38 *aak-2(gt33)* X.

The transgenic strains PHX893 (=DB06) *nish-1(syb767),* PHX117 *sybIs62* [*f13e9.1*::EGFP::3xFLAG, *unc-119*(+) + P*myo-2*::mCherry], and PHX116 *sybIs62* [*f13e9.1*::EGFP::3xFLAG, *unc-119*(+) + P*myo-2*::mCherry] were generated by SunyBiotech. In PHX893 (=DB06) *nish-1(syb767),* the 3rd exon, 4th exon, and part of 5th exon, was deleted and replaced by a splice site and stop codon that terminates transcription and resulted in 873 bp homozygous deletion in the *nish-1* gene. This strain was subsequently outcrossed four times against WT background using PCR screening at each stage, and the final outcrossed strain was confirmed by sequencing. The details of the primers are documented in Supplementary Table 2. PHX117 and PHX116 contain integrated full-length rescue of the *nish-1* locus within the fosmid generated in the WT background. Specifically, the integrated rescue contained the WRM0616C_F02(pRedFlp-Hgr)(F13E9.1[33366]::S0001_pR6K_Amp_2xTY1ce_EGFP_FR T_rpsl_neo_FRT_3xFlag)dFRT::unc-119-Nat) fosmid construct derived from the copy-number inducible vector pCC1Fos grown up in *E.coli* strain EPI300. Fosmid DNA was extracted and added to a 20 ng/µL DNA mixture containing both the purified fosmid and pCFJ90 P*myo-2*::mcherry. The solution was microinjected into 10-30 adult WT gonads to generate a line of extrachromosomal transgenic arrays. Stable lines were then generated by irradiating with a γ dose of 40 GRAY to produce two lines of the same genotype, PH117, and PH116. The previously produced, PHX893 (F13E9.1(*syb767*)IV) was then crossed 4 times with PX116 to generate the two rescue lines: PHX945 (=DB03) and PHX946 (=DB04) *f13e9.1 (syb767)* IV; *sybIs62* [*f13e9.1*::EGFP::3xFLAG, *unc-119*(+) + P*myo-2*::mCherry].

### Generation of UV-Killed OP50 E.coli

32 mL of unconcentrated live OP50 *E.coli* cultures were aliquoted to each 50 mL falcon tubes, mixed with absolute ethanol (8 mL) to yield a 20% ethanol solution as described in Calvert et al., 2016 and pipetted into T160 flasks to a volume of 150ml. Flasks were transferred to a UV-Linker machine (CL-1000 Ultraviolet crosslinker UVP) and irradiated for 120 minutes at 999,900 microjoules/cm^2^. 50ml aliquots of UV-killed *E.coli-*ethanol solution was pelleted at 3000 rpm for 10 minutes and resuspended to a 10X concentrate in 5 mL Lennox LB broth.

### Drug Treatment

All drug treatments, unless otherwise stated, were administered to *C.elegans* via the addition of the respective compound to NGM (Zheng et al. 2013). Aliquots of stock solutions of the drugs (rilmenidine and rapamycin) were prepared by dissolving them into DMSO solvent, such that final volume in NGM agar did not exceed 1%. NGM agar was cooled to below 65°C after same-day autoclaving, and respective drugs were concomitantly added. Plates were gently swirled to homogenize the solution and left to dry for 1 hour in a hood. 60 µl of 10X concentrated UV-killed OP50 *E.coli* was then spotted to the centre of the plate and again the plates were left to dry for 30 minutes in a hood. Unless otherwise stated, plates were prepared one day before use.

### Lifespan Assay

Well-fed gravid hermaphrodites were L1 synchronized overnight at 20°C, after hypochlorite treatment. The next day, L1s were transferred to NGM agar plates seeded with 10X OP50 *E.coli* and allowed to develop for 52 hours at 20°C until late L4/adult. At L4/adult molt, animals were transferred to 6-well lifespan assay plates, containing 3 ml of NGM per well with test compounds and 100 µg/ml 5-Fluoro-2’deoxyuridine (FUdR) (Calvert et al. 2016). Plates were maintained at 20°C in the dark, wrapped in parafilm to moderate humidity. Animals were counted every 2 to 3 days, and dead ones (failure to respond to no more than 3 “prods” with a platinum pick) were removed at each inspection. 60 μL of UV-killed OP50 was added three times a week to all plates until day 15 of the assay by which point low-food consumption negated the need for additional *E.coli* (Calvert et al. 2016).

For late administration lifespan assays, at day 12, animals were washed of lifespan assay plates with M9 buffer and carefully pipetted onto new drug plates prepared on day 11, seeded with UV-killed *E.coli*, and contained freshly prepared test compounds and 100 µg/ml FUdR. For RNAi lifespan analyses, L1 synchronized WT *C. elegans* were grown on live OP50 *E.coli* until late L4-stage. At L4 stage, animals were then transferred to NGM plates with 100 µM IPTG, 400 µM FUdR, and 50 µg/ml carbenicillin with either 200 µM rilmenidine or vehicle seeded with live *E.coli* (HT115) expressing an empty vector, *bec-1* dsRNA or *lgg-1* dsRNA. Plates were incubated for 1–2 days at room temperature prior to inducing dsRNA expression. Animals were maintained on the same drug plates for the entirety of their lifespan and scored for survival every 2-3 days.

For all lifespan assays, unless otherwise stated, at least 150 animals per genotype and/or condition were used across at least three independent trials. Animals that crawled into the agar, or experienced matricidal events were removed from the assay and not included in the life span calculations. The Kaplan-Meir nonparametric method was used for estimating survival curves. For each condition and/or genotype, survival results were pooled. Lifespan statistics were calculated using the OASIS2 online tool (Han et al., 2016). P-values relating to survival differences between populations were calculated using a log-rank (Mantel-Cox method) test.

### Automated Lifespan Measurements

Automated survival analysis was performed as described in (Statzer et al. 2020) employing the lifespan machine setup developed by Stroustrup and colleagues (Stroustrup et al. 2013). Briefly, C. elegans were age-synchronized using bleach lysis and kept in L1 culture (52h, 20°C), placed on live OP50 until L4 stage, and subsequently transferred to plates containing the drug, FUdR (50 µg/ml) and dead OP50 from L4 to day 4. Lastly, the animals were moved to tight-fitting Petri dishes (BD Falcon Petri Dishes, 50 x 9 mm) containing the drug, dead bacteria, and FUdR and imaged until the end of life.

Rilmenidine hemifumarate was dissolved in 1% DMSO and supplemented to the agar just before pouring to yield 200 µM, 300 µM, and 400 µM final concentrations. When using dead bacteria, the agar was additionally supplemented with Nystatin (44 U/ml) and Penicillin-Streptomycin (50 U/mL).

Air-cooled Epson V800 scanners were utilized for all experiments operating at a scanning frequency of one scan per 30 minutes. To limit condensation, the tight-fitting plates were dried without lids in a laminar flow hood for 40 minutes before starting the experiment.

Furthermore, temperature probes (Thermoworks, Utah, U.S.) were used to monitor the temperature on the scanner flatbed and maintain 20.0°C. Animals that left the imaging area during the experiment were censored. Automated lifespan results were validated using manual lifespan measurements as described in (Ewald et al. 2016) by picking L4 from normal culturing plates onto the corresponding assay plates. Lifespans were calculated from the L4 stage (= day 0).

### Body Size Phenotyping

Synchronized day 1 *C. elegans* hermaphrodites cultured on live OP50 *E. coli* NGM plates were measured for body length and width in brightfield at 10X objective on a Zeiss Axio Observer following paralysis in 20 mM tetramisole. Between 10-20 animals per data point were used. The animal’s length was measured from head (most visibly anterior buccal line) to a position where the tail tapered to a 10 ± 0.5 *μm* diameter (Petzold et al., 2011). Body width measurements were taken from the posterior vulval peak to the corresponding outer edge of the intestinal cuticle (Collins 2007). Measurements were made using the segmented lines function on Image J software normalized to the scale bar.

### Developmental Measurements

Bleach synchronized animals that had been halted in L1 starvation for 24h were measured for development following exposure to food source (Schindler et al. 2014). ∼ 500 L1s per genotype/condition were placed onto NGM plates containing rilmenidine and spotted with live OP50 *E.coli* and allowed to develop at 20°C. After 24 hours, measurement of development was conducted at different time points: 0 hours (24 hours exposure to food source), 3 hours, 6 hours, and then 24 hours (48 hours exposure to food source). Per trial and time point, 10 animals were immobilized and mounted onto glass slides in 20 mM solution of tetramisole hydrochloride and then imaged at a 10X objective for body length and 60X objective for vulval development in brightfield on a Zeiss Axio Observer. Vulval development was scored using visual identification of late larval stage vulval checkpoints detailed by Schindler (2014). Statistical significance of differences in vulval development was determined by Two-way ANOVA.

### Motility Assay

NGM agar plates were firmly tapped onto the microscope to stimulate movement. In responding animals, not impeded by OP50, body bends were counted for 30 seconds in 10 animals for each condition at each time point (5, 10, and 15 days post-L4 molt) per trial across three independent trials to a total of 30 animals per genotype and condition. Mean deterioration in motility per genotype and/or condition was compared by 2-factor repeated ANOVA and Tukey post hoc correction whilst individual time-point comparisons were tested by student’s *t*-test.

### Protein Extraction

∼500 day-1 adult animals were transferred to either empty NGM plates to induce starvation (+ve control) or to UV-killed OP50 *E.coli* NGM plates containing requisite drugs or vehicles. After 24 hours, animals were collected in an M9 buffer and centrifuged at 1000 rpm for 2 minutes to yield a wet pellet. Animals were then re-suspended on ice in 50 µl of RIPA buffer with added phosphatase inhibitors (Cell Signalling Technologies #9806) and complete EDTA-free protease inhibitor cocktail (Roche # 04693159001) before sonication: 10 sec ON/1 minute OFF, 14% amplitude 3 times until the vast majority of animals were dissolved. Lysates were then spun down twice at 13,000 g/ 20 minutes 4°C and immediately frozen at −20°C for no more than one week before use.

### Western Blot Analysis

Total protein lysate concentrations were quantified using a bicinchoninic acid (BCA) assay (BioRAD) according to the manufacturer’s protocol.

Protein lysates were heated for 5 minutes at 95°C, cooled on ice for 2 minutes, and pelleted for 3 minutes at 13,000 g. 25 µg of protein was loaded per well into an 18 well 10% TGX™ Precast Gel (BioRad), submerged in running buffer (25 mM Tris, 192 mM glycine, 0.1% SDS, pH 8.3) and separated at 300V for approximately 30 minutes. Gels were transferred via Trans-Blot® Turbo™ Transfer System to nitrocellulose membranes and blocked for 60 minutes in blocking buffer (Licor Odyssey® Blocking Buffer #P/N 927-40100). Blots were incubated overnight with the primary antibody Phospho-ERK (Cell Signalling Technology, Catalogue number: 9101) at 1:1000 in blocking buffer (Licor Odyssey® Blocking Buffer #P/N 927-40100) (Gee et al., 2013) with gentle rocking. The following morning, blots were washed three times in TBST before being incubated for 1 hour with the secondary antibody IRDye® 800CW Goat anti-Rabbit IgG at a concentration of 1:10,00 in blocking buffer w/ 0.1% tween. α-tubulin was used as the loading control [Primary anti-α-tubulin : ab72910 and Secondary antibody: IRDye® 680RD Goat anti-Mouse IgG]. Bound antibody was detected using Odyssey® CLx Imaging System. Changes in ERK phosphorylation are often calculated using phospho-ERK:Total ERK ratios (Kao et al. 2004; Nykamp et al. 2008). However, given the frequent use of ERK as a loading control, we calculated changes in phospho-ERK via the surrogate loading control α-tubulin.

For all immunoblotting, three independent trials for each genotype and/or condition were completed and mean densitometric ratios between loading control and pERK minus local median background were calculated using Image Studio Lite as per software guidelines (Licor Biosciences 2013). Results were considered significant if the magnitude of the %change exceeded the CV by at least x1.5.

### Thermotolerance and Recovery Assay

Well-fed gravid hermaphrodites were bleach synchronised and resultant embryos were allowed to grow for 52 hours at 20°C on NGM plates seeded with live OP50 *E.coli*. At late-L4 stage, animals were transferred to seeded UV-killed OP50 *E.coli* NGM plates containing 200-400 µM rilmenidine, or vehicle for 24h. After 24h of the respective drug exposure, plates were upshifted to an incubator preset to 37°C to induce heat shock. After 3 hours, plates were moved to 20°C and scored for survival after a ∼20h “recovery period” (Kumsta et al. 2017). At least 100 animals per condition across three identically designed independent trials were scored as either dead or alive by their ability to respond to no more than 3 “prods” with a platinum pick. Percentage survival was calculated and populations were compared by one-way ANOVA of variance with Tukey post hoc correction.

### Measurement of Autophagy

Gravid hermaphrodites were bleach synchronized and resultant embryos were allowed to grow for 52 hours at 20°C on NGM plates seeded with live OP50 *E.coli*. At late-L4 stage, animals were transferred to either empty NGM plates to induce starvation (+ve control) or to UV-killed OP50 *E.coli* NGM plates containing requisite drugs or vehicle (1% DMSO). Animals were then incubated at 20°C for 24 hours to maximize drug absorption and efficacy (Zheng et al., 2013). After 24 hours, approximately 10 day-1 adults per condition were imaged at a 10X objective on a Zeiss Axio Observer using a 150 ms exposure time. In total, at least 25 images across three independent trials were collected and pooled for each condition (Eisenberg et al. 2009; Morselli et al. 2010). Levels of autophagy induction were quantified using Zen software. Three 1 mm^2^ boxes were sequentially assigned to the most fluorescent areas of the posterior intestine for each animal and with set histogramic parameters (black: 0 gamma:1.0 white: 1000 or 5000 for highly fluorescent animals), the total number of visible puncta were manually counted in each box, and a mean number of puncta calculated per mm^2^ of animal intestine. Median number of puncta per mm^2^ posterior intestine for all animals in each condition was calculated and compared using one-factor (ANOVA) variance analysis corrected by the post hoc Bonferroni test.

### PolyQ Protein Aggregation Assay

Gravid hermaphrodites were bleached, and synchronized L1s were transferred to NGM plates containing rilmenidine or DMSO. After incubation for the indicated periods of time, approximately 30 animals per condition across three trials were imaged under a Zeiss Axio Observer microscope at 10X objective. The number of polyQ40::YFP aggregates in body wall muscle was counted using ZenBlue software with set histogramic parameters (black: 500 gamma:1.0 white: 5000). Approximately 50 animals across three independent trials were randomly selected for each treatment group and scored for the number of aggregates. Groups were compared by two-factor repeated measures ANOVA and Tukey posthoc correction.

### RNA-sequencing Data Processing

RNAseq data of gene expression changes induced by rilmenidine in mouse liver after 1 month of oral administration was obtained from (Tyshkovskiy et al. 2019). In addition, we performed RNA sequencing of corresponding kidney samples from 8 control mice and 4 mice subjected to the drug for 1 month. RNA was extracted from tissues with the PureLink RNA Mini Kit as described in the protocol. Libraries were prepared and sequenced with 100 bp read length option on the Illumina HiSeq 2500.

Quality filtering and adapter removal were performed using Trimmomatic (version 0.32). Processed/cleaned reads were then mapped with STAR (version 2.5.2b) and counted via featureCounts (version 1.5). To filter out genes with low number of reads, we left only genes with at least 6 reads in at least 66.6% of the samples separately for each tissue. Filtered data was then passed to RLE normalization (Anders & Huber 2010). Differentially expressed genes between control and treated mice were identified using edgeR. For each gene, we calculated the p-value of its logFC in response to rilmenidine compared to control independently for every tissue. We then converted them to log_10_(p-value) corrected by the sign of regulation, calculated as:

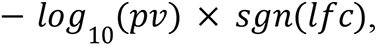

where *pv* and *lfc* are p-value and logFC of a certain gene, respectively, and *sgn* is signum function (is equal to 1, −1 and 0 if the value is positive, negative and zero, respectively). The resulting pre-ranked list of genes was used to identify significant associations with the biomarkers of aging, lifespan extension and specific intracellular functions.

### *In-silico* Association Analysis

To identify functions enriched by genes perturbed by rilmenidine, we performed gene set enrichment analysis (GSEA) (Subramanian et al. 2005) on a pre-ranked list of genes, obtained as described earlier. REACTOME, KEGG, and GO BP from Molecular Signature Database (MSigDB) have been used as gene sets for GSEA. Only functions related to the processes identified in *C. elegans* (insulin, FOXO, ERK and mTOR signaling, autophagy and proteolysis) were chosen for the analysis.

To identify associations between rilmenidine’s effect in mice and biomarkers of lifespan-extending interventions and aging, we employed a GSEA-based algorithm developed in our previous study (Tyshkovskiy et al. 2019). First, for every trait, we specified significant genes, using the FDR threshold of 0.05. Among the remaining genes, we selected 1000 genes with the highest absolute logFC and divided them into up- and downregulated genes. These lists were considered as gene sets. Signatures of lifespan-extending interventions were taken from (Tyshkovskiy et al. 2019), while signatures of aging in liver and kidney were identified from GSE132040 dataset derived by Tabula Muris Consortium (Schaum et al. 2020). Then we calculated normalized enrichment scores (NES) separately for the up- and downregulated lists of each gene set as described in (Lamb 2006) and defined the final NES as a mean of the two. To calculate the statistical significance of the NES, we performed a permutation test where we randomly selected gene sets of the same size. p-value of association was calculated as the frequency of observed final NES being bigger by absolute value than random final NES obtained from 5,000 random permutations. The resulting p-values were further adjusted for multiple hypothesis testing using Benjamini-Hochberg (BH) procedure (Benjamini & Hochberg 1995). Association was considered significant if the adjusted p-value was smaller than 0.1. To visualize the results of the association analysis, we converted adjusted p-values into significance scores as:

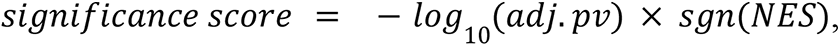

where adj.pv and NES are BH adjusted p-value and final NES, respectively.

## Author contributions

All authors participated in analyzing and interpreting the data. DFB, JPdM, and CYE designed the experiments. DFB and CYE performed manual lifespan assays, and CS performed automated lifespan machine assays. AT and VNG performed mice experiments and analyses. All other experiments were performed by DFB. AG, CWB, CYE, and JPdM wrote the manuscript in consultation with the other authors.

## Author Information

JPM is CSO of Centaura, a company that aims to prevent and reverse aging, an advisor for the Longevity Vision Fund, NOVOS and YouthBio as well as the founder of Magellan Science Ltd, a company providing consulting services in longevity science. CYE is a co-founder of AVEA Life AG and an advisor for Maximon AG Longevity Startup builder. The other authors have no competing interests to declare. Correspondence should be addressed to C. Y. E. and J.P.d.M.

## Acknowledgment

We thank Dominic Alcock and Alan Morgan for assistance and advice with the worm experiments. Further thanks to the de Magalhães and Ewald labs for critical feedback on the manuscript and to Wormbase for curating and updating the *nish-1* gene information. Some strains were provided by the CGC, which is funded by NIH Office of Research Infrastructure Programs (P40 OD010440). Graphic illustrations in Figure 2D and 6F were created with BioRender.com (PQ22YK2PVV and LT22YK2745). Funding from the Swiss National Science Foundation 163898 and 190072 to CS, AG, and CYE, and from LongeCity and the Biotechnology and Biological Sciences Research Council (BB/R014949/1) to JPM.

## References

Admasu TD, Chaithanya Batchu K, Barardo D, Ng LF, Lam VYM, Xiao L & Gruber J (2018) Drug Synergy Slows Aging and Improves Healthspan through IGF and SREBP Lipid Signaling, Dev Cell, 47(1), 67–79 e65.

Alahari SK, Lee JW & Juliano RL (2000) Nischarin, a Novel Protein That Interacts with the Integrin α5 Subunit and Inhibits Cell Migration. Available at: https://doi.org/10.1083/jcb.151.6.1141.

Alonso N, Diaz Nebreda A, Monczor F, Gutkind JS, Davio C, Fernandez N & Shayo C (2016) PI3K pathway is involved in ERK signaling cascade activation by histamine H2R agonist in HEK293T cells. Available at: https://doi.org/10.1016/j.bbagen.2016.06.016.

Anders S & Huber W (2010) Differential expression analysis for sequence count data. Genome Biol. 11, R106.

Benjamini Y & Hochberg Y (1995) Controlling the False Discovery Rate: A Practical and Powerful Approach to Multiple Testing. J. R. Stat. Soc. Ser. B Methodol. 57, 289–300.

Berger Z, Ravikumar B, Menzies FM, Oroz LG, Underwood BR, Pangalos MN, Schmitt I, Wullner U, Evert BO, O’Kane CJ & Rubinsztein DC (2006) Rapamycin alleviates toxicity of different aggregate-prone proteins. Available at: https://doi.org/10.1093/hmg/ddi458.

Blackwell TK, Sewell AK, Wu Z & Han M (2019) TOR Signaling in Caenorhabditis elegans Development, Metabolism, and Aging. Genetics 213, 329–360.

Blackwell TK, Steinbaugh MJ, Hourihan JM, Ewald CY & Isik M (2015) SKN-1/Nrf, stress responses, and aging in Caenorhabditis elegans. Free Radic. Biol. Med. 88, 290–301.

Calvert S, Tacutu R, Sharifi S, Teixeira R, Ghosh P & de Magalhaes JP (2016) A network pharmacology approach reveals new candidate caloric restriction mimetics in C, elegans. Aging Cell, 15(2), 256–266.

Chung KW & Chung HY (2019) The effects of calorie restriction on autophagy: Role on aging intervention. Available at: https://doi.org/10.3390/nu11122923.

Collins TJ (2007) ImageJ for microscopy. BioTechniques 43, S25–S30.

Crompton M, Purnell T, Tyrer HE, Parker A, Ball G, Hardisty-Hughes RE, Gale R, Williams D, Dean CH, Simon MM, Mallon AM, Wells S, Bhutta MF, Burton MJ, Tateossian H & Brown SDM (2017) A mutation in Nischarin causes otitis media via LIMK1 and NF-κB pathways. Available at: https://doi.org/10.1371/journal.pgen.1006969.

Dardonville C & Rozas I (2004) Imidazoline binding sites and their ligands: an overview of the different chemical structures, Med Res Rev, 24(5), 639–661.

Dong S, Baranwal S, Garcia A, Serrano-Gomez SJ, Eastlack S, Iwakuma T, Mercante D, Mauvais-Jarvis F & Alahari SK (2017) Nischarin inhibition alters energy metabolism by activating AMP-activated protein kinase. Available at: https://doi.org/10.1074/jbc.m117.784256.

Dontenwill M, Pascal G, Piletz JE, Chen M, Baldwin J, Rondé P, Dupuy L, Urosevic D, Greney H, Takeda K & Bousquet P (2003) IRAS, the human homologue of Nischarin, prolongs survival of transfected PC12 cells [3]. Available at: https://doi.org/10.1038/sj.cdd.4401275.

Eisenberg T, Knauer H, Schauer A, Büttner S, Ruckenstuhl C, Carmona-Gutierrez D, Ring J, Schroeder S, Magnes C, Antonacci L, Fussi H, Deszcz L, Hartl R, Schraml E, Criollo A, Megalou E, Weiskopf D, Laun P, Heeren G, Breitenbach M, Grubeck-Loebenstein B, Herker E, Fahrenkrog B, Fröhlich K-U, Sinner F, Tavernarakis N, Minois N, Kroemer G & Madeo F (2009) Induction of autophagy by spermidine promotes longevity. Nat. Cell Biol. 11, 1305–1314.

Espada L, Dakhovnik A, Chaudhari P, Martirosyan A, Miek L, Poliezhaieva T, Schaub Y, Nair A, Döring N, Rahnis N, Werz O, Koeberle A, Kirkpatrick J, Ori A & Ermolaeva MA (2020) Loss of metabolic plasticity underlies metformin toxicity in aged Caenorhabditis elegans. Available at: https://doi.org/10.1038/s42255-020-00307-1.

Ewald CY, Landis JN, Porter Abate J, Murphy CT & Blackwell TK (2015) Dauer-independent insulin/IGF-1-signalling implicates collagen remodelling in longevity. Nature 519, 97–101.

Ewald CY, Marfil V & Li C (2016) Alzheimer-related protein APL-1 modulates lifespan through heterochronic gene regulation in Caenorhabditis elegans. Aging Cell 15, 1051–1062.

Farrelly C (2010) Why aging research? The moral imperative to retard human aging. Ann. N. Y. Acad. Sci. 1197, 1–8.

Fontana L, Kennedy BK, Longo VD, Seals D & Melov S (2014) Medical research: treat ageing. Nature 511, 405–407.

Fontana L, Partridge L & Longo VD (2010) Extending Healthy Life Span--From Yeast to Humans. Available at: https://doi.org/10.1126/science.1172539.

Goldman DP, Cutler D, Rowe JW, Michaud P-C, Sullivan J, Peneva D & Olshansky SJ (2013) Substantial health and economic returns from delayed aging may warrant a new focus for medical research. Health Aff. Proj. Hope 32, 1698–1705.

Gosai SJ, Kwak JH, Luke CJ, Long OS, King DE, Kovatch KJ, Johnston PA, Shun TY, Lazo JS, Perlmutter DH, Silverman GA & Pak SC (2010) Automated High-Content Live Animal Drug Screening UsingC. Available at: https://doi.org/10.1371/journal.pone.0015460.

Greer EL, Dowlatshahi D, Banko MR, Villen J, Hoang K, Blanchard D, Gygi SP & Brunet A (2007) An AMPK-FOXO Pathway Mediates Longevity Induced by a Novel Method of Dietary Restriction in C. Available at: https://doi.org/10.1016/j.cub.2007.08.047.

Hansen M, Chandra A, Mitic LL, Onken B, Driscoll M & Kenyon C (2008) A Role for Autophagy in the Extension of Lifespan by Dietary Restriction in C. elegans S. K. Kim, ed. PLoS Genet. 4, e24.

Hansen M, Taubert S, Crawford D, Libina N, Lee SJ & Kenyon C (2007) Lifespan extension by conditions that inhibit translation in Caenorhabditis elegans. Available at: https://doi.org/10.1111/j.1474-9726.2006.00267.x.

Hosono R, Sato Y, Aizawa SI & Mitsui Y (1980) Age-dependent changes in mobility and separation of the nematode Caenorhabditis elegans. Available at: https://doi.org/10.1016/0531-5565(80)90032-7.

Hsu AL, Murphy CT & Kenyon C (2003) Regulation of aging and age-related disease by DAF-16 and heat-shock factor. Available at: https://doi.org/10.1126/science.1083701.

Ingram DK & Roth GS (2015) Calorie restriction mimetics: can you have your cake and eat it, too? Ageing Res Rev, 20, 46-62,

Jarzebski MP, Elmqvist T, Gasparatos A, Fukushi K, Eckersten S, Haase D & Pu J (2021) Ageing and population shrinking: implications for sustainability in the urban century, npj Urban Sustainability, 1(1), 17.

Kahn LP & Nolan JV (2000) Kinetics of allantoin metabolism in sheep, Br J Nutr, 84(5), 629–634.

Kao G, Tuck S, Baillie D & Sundaram MV (2004) C. elegans SUR-6/PR55 cooperates with LET-92/protein phosphatase 2A and promotes Raf activity independently of inhibitory Akt phosphorylation sites. Development 131, 755–765.

Kumsta C, Chang JT, Schmalz J & Hansen M (2017) Hormetic heat stress and HSF-1 induce autophagy to improve survival and proteostasis in C. Available at: https://doi.org/10.1038/ncomms14337.

Lakowski B & Hekimi S (1998) The genetics of caloric restriction in Caenorhabditis elegans. Available at: https://doi.org/10.1073/pnas.95.22.13091.

Lamb J (2006) The Connectivity Map: Using Gene-Expression Signatures to Connect Small Molecules, Genes, and Disease. Science 313, 1929–1935.

Li J, Pak SC, O’Reilly LP, Benson JA, Wang Y, Hidvegi T, Hale P, Dippold C, Ewing M, Silverman GA & Perlmutter DH (2014) Fluphenazine Reduces Proteotoxicity in C. Available at: https://doi.org/10.1371/journal.pone.0087260.

Liang Y, Liu C, Lu M, Dong Q, Wang Z, Wang Z & Wang Z (2018) Calorie restriction is the most reasonable anti-ageing intervention: a meta-analysis of survival curves, Sci Rep, 8(1), 5779.

Licor Biosciences (2013) Tutorial Guide Featuring Image Studio Analysis Software Version 3.1 CLx.

López-Otín C, Blasco MA, Partridge L, Serrano M & Kroemer G (2013) The hallmarks of aging. Cell 153, 1194–1217.

Luca ND, Izzo R, Fontana D, Iovino G, Argenziano L, Vecchione C & Trimarco B (2000) Haemodynamic and metabolic effects of rilmenidine in hypertensive patients with metabolic syndrome X. Available at: https://doi.org/10.1097/00004872-200018100-00021.

Ludewig AH, Gimond C, Judkins JC, Thornton S, Pulido DC, Micikas RJ, Döring F, Antebi A, Braendle C & Schroeder FC (2017) Larval crowding accelerates C. Available at: https://doi.org/10.1371/journal.pgen.1006717.

Madeo F, Zimmermann A, Maiuri MC & Kroemer G (2015) Essential role for autophagy in life span extension. Available at: https://doi.org/10.1172/jci73946.

de Magalhães JP, Wuttke D, Wood SH, Plank M & Vora C (2012) Genome-environment interactions that modulate aging: powerful targets for drug discovery. Pharmacol. Rev. 64, 88–101.

Mariño G, Pietrocola F, Madeo F & Kroemer G (2014) Caloric restriction mimetics: Natural/physiological pharmacological autophagy inducers. Available at: https://doi.org/10.4161/auto.36413.

Matai L, Sarkar GC, Chamoli M, Malik Y, Kumar SS, Rautela U, Jana NR, Chakraborty K & Mukhopadhyay A (2019) Dietary restriction improves proteostasis and increases life span through endoplasmic reticulum hormesis. Available at: https://doi.org/10.1073/pnas.1900055116.

Morselli E, Maiuri MC, Markaki M, Megalou E, Pasparaki A, Palikaras K, Criollo A, Galluzzi L, Malik SA, Vitale I, Michaud M, Madeo F, Tavernarakis N & Kroemer G (2010) Caloric restriction and resveratrol promote longevity through the Sirtuin-1-dependent induction of autophagy. Cell Death Dis. 1, e10–e10.

Most J, Tosti V, Redman LM & Fontana L (2017) Calorie restriction in humans: An update, Ageing Res Rev, 39, 36–45.

Nykamp K, Lee M-H & Kimble J (2008) C. elegans La-related protein, LARP-1, localizes to germline P bodies and attenuates Ras-MAPK signaling during oogenesis. RNA 14, 1378–1389.

Onken B & Driscoll M (2010) Metformin Induces a Dietary Restriction-Like State and the Oxidative Stress Response to Extend C. Available at: https://doi.org/10.1371/journal.pone.0008758.

Perera ND, Sheean RK, Lau CL, Shin YS, Beart PM, Horne MK & Turner BJ (2017) Rilmenidine promotes MTOR-independent autophagy in the mutant SOD1 mouse model of amyotrophic lateral sclerosis without slowing disease progression. Available at: https://doi.org/10.1080/15548627.2017.1385674.

Perera ND, Sheean RK, Lau CL, Shin YS, Beart PM, Horne MK & Turner BJ (2018) Rilmenidine promotes MTOR-independent autophagy in the mutant SOD1 mouse model of amyotrophic lateral sclerosis without slowing disease progression. Available at: https://doi.org/10.1080/15548627.2017.1385674.

Perera ND, Tomas D, Wanniarachchillage N, Cuic B, Luikinga SJ, Rytova V & Turner BJ (2021) Stimulation of mTOR-independent autophagy and mitophagy by rilmenidine exacerbates the phenotype of transgenic TDP-43 mice. Available at: https://doi.org/10.1016/j.nbd.2021.105359.

Piletz J (2003) Cell Signaling by Imidazoline-1 Receptor Candidate, IRAS, and the Nischarin Homologue. Available at: https://doi.org/10.1196/annals.1304.053.

Public Health England (2020) Public Health England. In Hypertension prevalence estimates in England, 2017, *London*: . *PHE publications*.

Ravikumar B, Duden R & Rubinsztein DC (2002) Aggregate-prone proteins with polyglutamine and polyalanine expansions are degraded by autophagy. Available at: https://doi.org/10.1093/hmg/11.9.1107.

Ravikumar B, Vacher C, Berger Z, Davies JE, Luo S, Oroz LG, Scaravilli F, Easton DF, Duden R, O’Kane CJ & Rubinsztein DC (2004) Inhibition of mTOR induces autophagy and reduces toxicity of polyglutamine expansions in fly and mouse models of Huntington disease. Available at: https://doi.org/10.1038/ng1362.

Reid JL (2001) Update on rilmenidine: Clinical benefits. Available at: https://doi.org/10.1016/s0895-7061(01)02239-7.

Robida-Stubbs S, Glover-Cutter K, Lamming D, Mizunuma M, Narasimhan S, Neumann-Haefelin E, Sabatini D & Blackwell T (2012) TOR Signaling and Rapamycin Influence Longevity by Regulating SKN-1/Nrf and DAF-16/FoxO. Available at: https://doi.org/10.1016/j.cmet.2012.04.007.

Rose C, Menzies FM, Renna M, Acevedo-Arozena A, Corrochano S, Sadiq O, Brown SD & Rubinsztein DC (2010) Rilmenidine attenuates toxicity of polyglutamine expansions in a mouse model of Huntington’s disease. Available at: https://doi.org/10.1093/hmg/ddq093.

Sanchis A, García-Gimeno MA, Cañada-Martínez AJ, Sequedo MD, Millán JM, Sanz P & Vázquez-Manrique RP (2019) Metformin treatment reduces motor and neuropsychiatric phenotypes in the zQ175 mouse model of Huntington disease. Available at: https://doi.org/10.1038/s12276-019-0264-9.

Schaum N, Lehallier B, Hahn O, Pálovics R, Hosseinzadeh S, Lee SE, Sit R, Lee DP, Losada PM, Zardeneta ME, Fehlmann T, Webber JT, McGeever A, Calcuttawala K, Zhang H, Berdnik D, Mathur V, Tan W, Zee A, Tan M, The Tabula Muris Consortium, Almanzar N, Antony J, Baghel AS, Bakerman I, Bansal I, Barres BA, Beachy PA, Berdnik D, Bilen B, Brownfield D, Cain C, Chan CKF, Chen MB, Clarke MF, Conley SD, Darmanis S, Demers A, Demir K, de Morree A, Divita T, du Bois H, Ebadi H, Espinoza FH, Fish M, Gan Q, George BM, Gillich A, Gòmez-Sjöberg R, Green F, Genetiano G, Gu X, Gulati GS, Hahn O, Haney MS, Hang Y, Harris L, He M, Hosseinzadeh S, Huang A, Huang KC, Iram T, Isobe T, Ives F, Jones R, Kao KS, Karkanias J, Karnam G, Keller A, Kershner AM, Khoury N, Kim SK, Kiss BM, Kong W, Krasnow MA, Kumar ME, Kuo CS, Y. Lam J, Lee DP, Lee SE, Lehallier B, Leventhal O, Li G, Li Q, Liu L, Lo A, Lu W-J, Lugo-Fagundo MF, Manjunath A, May AP, Maynard A, McGeever A, McKay M, McNerney MW, Merrill B, Metzger RJ, Mignardi M, Min D, Nabhan AN, Neff NF, Ng KM, Nguyen PK, Noh J, Nusse R, Pálovics R, Patkar R, Peng WC, Penland L, Pisco AO, Pollard K, Puccinelli R, Qi Z, Quake SR, Rando TA, Rulifson EJ, Schaum N, Segal JM, Sikandar SS, Sinha R, Sit RV, Sonnenburg J, Staehli D, Szade K, Tan M, Tan W, Tato C, Tellez K, Dulgeroff LBT, Travaglini KJ, Tropini C, Tsui M, Waldburger L, Wang BM, van Weele LJ, Weinberg K, Weissman IL, Wosczyna MN, Wu SM, Wyss-Coray T, Xiang J, Xue S, Yamauchi KA, Yang AC, Yerra LP, Youngyunpipatkul J, Yu B, Zanini F, Zardeneta ME, Zee A, Zhao C, Zhang F, Zhang H, Zhang MJ, Zhou L, Zou J, Pisco AO, Karkanias J, Neff NF, Keller A, Darmanis S, Quake SR & Wyss-Coray T (2020) Ageing hallmarks exhibit organ-specific temporal signatures. Nature 583, 596–602.

Schindler AJ, Baugh LR & Sherwood DR (2014) Identification of Late Larval Stage Developmental Checkpoints in Caenorhabditis elegans Regulated by Insulin/IGF and Steroid Hormone Signaling Pathways K. Ashrafi, ed. PLoS Genet. 10, e1004426.

Shintani H, Shintani T, Ashida H & Sato M (2018) Calorie restriction mimetics: Upstream-type compounds for modulating glucose metabolism. Available at: https://doi.org/10.3390/nu10121821.

Statzer C, Jongsma E, Liu SX, Dakhovnik A, Wandrey F, Mozharovskyi P, Zülli F & Ewald CY (2021) Youthful and age-related matreotypes predict drugs promoting longevity. *Aging Cell*, e13441.

Statzer C, Meng J, Venz R, Bland M, Robida-Stubbs S, Patel K, Petrovic D, Emsley R, Liu P, Morantte I, Haynes C, Mair WB, Longchamp A, Filipovic M, Blackwell TK & Ewald CY (2020) *ATF-4 and hydrogen sulfide signalling mediate longevity from inhibition of translation or mTORC1*, Physiology. Available at: http://biorxiv.org/lookup/doi/10.1101/2020.11.02.364703 [Accessed September 4, 2021].

Stiernagle T (2006) Maintenance of C. elegans. *WormBook*. Available at: http://www.wormbook.org/chapters/www_strainmaintain/strainmaintain.html [Accessed September 4, 2021].

Stroustrup N, Ulmschneider BE, Nash ZM, López-Moyado IF, Apfeld J & Fontana W (2013) The Caenorhabditis elegans Lifespan Machine. Nat. Methods 10, 665–670.

Subramanian A, Tamayo P, Mootha VK, Mukherjee S, Ebert BL, Gillette MA, Paulovich A, Pomeroy SL, Golub TR, Lander ES & Mesirov JP (2005) Gene set enrichment analysis: A knowledge-based approach for interpreting genome-wide expression profiles. Proc. Natl. Acad. Sci. 102, 15545–15550.

Sun Z, Chang CH & Ernsberger P (2007) Identification of IRAS/Nischarin as an I1-imidazoline receptor in PC12 rat pheochromocytoma cells. Available at: https://doi.org/10.1111/j.1471-4159.2006.04413.x.

Sunayama J, Matsuda KI, Sato A, Tachibana K, Suzuki K, Narita Y, Shibui S, Sakurada K, Kayama T, Tomiyama A & Kitanaka C (2010) Crosstalk between the PI3K/mTOR and MEK/ERK pathways involved in the maintenance of self-renewal and tumorigenicity of glioblastoma stem-like cells. Available at: https://doi.org/10.1002/stem.521.

Tyshkovskiy A, Bozaykut P, Borodinova AA, Gerashchenko MV, Ables GP, Garratt M, Khaitovich P, Clish CB, Miller RA & Gladyshev VN (2019) Identification and Application of Gene Expression Signatures Associated with Lifespan Extension. Available at: https://doi.org/10.1016/j.cmet.2019.06.018.

Yang Y & Zhang L (2020) The effects of caloric restriction and its mimetics in Alzheimer’s disease through autophagy pathways. Available at: https://doi.org/10.1039/c9fo02611h.

You YJ, Kim J, Cobb M & Avery L (2006) Starvation activates MAP kinase through the muscarinic acetylcholine pathway in Caenorhabditis elegans pharynx. Available at: https://doi.org/10.1016/j.cmet.2006.02.012.

Zhang H, Chang JT, Guo B, Hansen M, Jia K, Kovács AL, Kumsta C, Lapierre LR, Legouis R, Lin L, Lu Q, Meléndez A, O’Rourke EJ, Sato K, Sato M, Wang X & Wu F (2015) Guidelines for monitoring autophagy in Caenorhabditis elegans. Available at: https://doi.org/10.1080/15548627.2014.1003478.

Zhang J & Abdel-Rahman AA (2005) Mitogen-Activated Protein Kinase Phosphorylation in the Rostral Ventrolateral Medulla Plays a Key Role in Imidazoline (I1)-Receptor-Mediated Hypotension. Available at: https://doi.org/10.1124/jpet.105.087510.

Zhang J & Abdel-Rahman AA (2006) Nischarin as a functional imidazoline (I1) receptor. Available at: https://doi.org/10.1016/j.febslet.2006.04.058.

Zhang L, Zhao TY, Hou N, Teng Y, Cheng X, Wang B, Chen Y, Jiang L, Wu N, Su RB, Yang X & Li J (2013) Generation and Primary Phenotypes of Imidazoline Receptor Antisera-Selected (IRAS) Knockout Mice. Available at: https://doi.org/10.1111/cns.12192.

Zhang Y, Lanjuin A, Chowdhury SR, Mistry M, Silva-García CG, Weir HJ, Lee CL, Escoubas CC, Tabakovic E & Mair WB (2019) Neuronal TORC1 modulates longevity via AMPK and cell nonautonomous regulation of mitochondrial dynamics in C. Available at: https://doi.org/10.7554/elife.49158.

Zheng S-Q, Ding A-J, Li G-P, Wu G-S & Luo H-R (2013) Drug absorption efficiency in Caenorhbditis elegans delivered by different methods. PloS One 8, e56877.

Zhou KI, Pincus Z & Slack FJ (2011) Longevity and stress in Caenorhabditis elegans. Available at: https://doi.org/10.18632/aging.100367.

Zhu X, Shen W, Liu Z, Sheng S, Xiong W, He R, Zhang X, Ma L & Ju Z (2021) Effect of Metformin on Cardiac Metabolism and Longevity in Aged Female Mice. Available at: https://doi.org/10.3389/fcell.2020.626011.

